# Symbolic Information Flow Measurement (SIFM): A Software for Measurement of Information Flow Using Symbolic Analysis

**DOI:** 10.1101/785782

**Authors:** Dhurata Nebiu, Hiqmet Kamberaj

## Abstract

Symbolic Information Flow Measurement software is used to compute the information flow between different components of a dynamical system or different dynamical systems using symbolic transfer entropy. Here, the time series represents the time evolution trajectory of a component of the dynamical system. Different methods are used to perform a symbolic analysis of the time series based on the coarse-graining approach by computing the so-called embedding parameters. Information flow is measured in terms of the so-called average symbolic transfer entropy and local symbolic transfer entropy. Besides, a new measure of mutual information is introduced based on the symbolic analysis, called symbolic mutual information.

## 1 Introduction

In many fields of research, we often need to determine the causal directions between parts of the same system or coupled systems. That is because we have to understand system dynamics and make an estimation of its actual physical structure. The process includes observation of the system, recording its behavior as a trajectory in phase space, or the so-called here time series of signals and analyzing this time series. The simplest measure of statistical dependency is the Pearson correlation coefficient [1]. However, it does not imply causality, and also, it detects only the linear correlations and omits the non-linear correlations, and furthermore, it is not sensitive to fluctuations in perpendicular directions, but only to those that distribute along with co-linear directions [2] (and the references therein). The fundamental concept for the dependence of one variable *Y* measured over time on another variable *X* measured synchronously is the Granger causality [3]. While Granger defined the direction of interaction in terms of the contribution of *X* in predicting *Y*, many variations of this concept have been developed, starting with linear approaches in the time and frequency domain.

The information theory measure of transfer entropy quantifies the statistical coherence between two processes that evolve in time. Transfer entropy (TE) is introduced by Schreiber [4] as the deviation from the independence of the state transition (from the previous state to the next state) of an information destination *X* from the (previous)state of an information source *Y* TE is asymmetric, and it is a measure of the information flow between dynamical variables, which correspond to either different components of a dynamical system or different dynamical systems. As an asymmetric measure, TE can distinguish between the dynamical macro-variable characterizing the actual dynamic behavior of the physical system (*source*), and the other macro-variable that responds to the changes (*sink*). [4]

Computing TE is a challenging problem due to its computational complexity. Therefore, different numerical recipes have been suggested [5]. TE has already been used for time series analysis in different fields, such as clinical electroencephalography [4, 6, 7], financial data [8], and biophysics [2]. Many algorithms used to calculate TE are subject to statistical errors, and moreover, reliable estimations of the transfer entropy are data intensive. [2]

In this work, we present a software, written in Fortran 90 programming language, used to perform symbolic information flow measurements.

## 2 Problems and Background

### Transfer Entropy

A dynamical system or a component of it can be characterized by the dynamical variable *X*, which is a time-dependent variable. Usually, the dynamical process is a *time-discrete process*, with time as t_k_ = kΔ t, where *k* is an integer and Δ*t* is time resolution of the process, and hence, X(t) = {x(t)}_k=0,…, N-1_, where N is the total number of time frames. Here, we prefer to use the notation X(t_k_) ≡ x_k_.

The dynamics of a time-discrete process can be characterized by the method of time-delayed embedding [9, 10, 11, 12], in which a so-called *state vector* in a *m*-dimensional space of a discrete process X(t) is determined as:

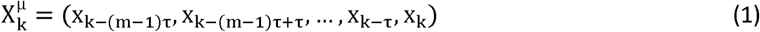

where τ is the time lag, which, in general, is a multiple of Δt, and k = (m− 1) τ,..,N −1. Also, the following vector is introduced, 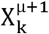

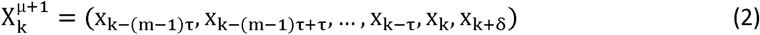

where X (t_k+δ_) ≡ x _k+δ_

It is interesting to note that both m and τ are characteristics of each time series, and furthermore, their choice is crucial in reconstructing the dynamical structure of the process in its higher dimension space, as followed by our discussion according to the literature [2]. For clarity, we denote by *μ* ≡ (*m, τ*), *μ*+1 ≡ (*m*+ 1, *τ*), and *k*_0_≡ max_*x,y*_((*m* − 1) *τ*).

Now, consider two-time series, namely 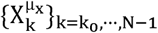 and 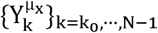, then the transfer entropy can be defined as [4] (using the notations as in the Ref. [2])

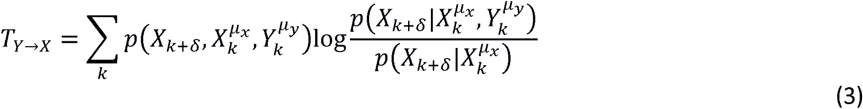

where *k* is a time index, 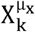 and 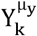 represent *m*_*x*_ and *m*_*y*_ past values of X and Y processes up to and including time step *K* defined according to Eq. (1). Here,*m*_*x*/*y*_ and *τ*_*x*/*y*_ represent the embedded dimension and time lag of each process. In this formulation, the transfer entropy represents a dynamical measure, as a generalization of the entropy rate to more than one element to form a mutual information rate (or conditional mutual information) [4].

In Eq. (3), 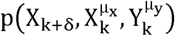 is the joint probability distribution of observing the future value X_k+ δ_ and the histories of 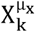 and 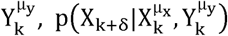 is the conditional probability of observing the future value X_k+δ_ given the past values of both processes 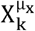 and 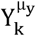, and 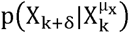 is the conditional probability of observing the future X_k+ δ_ knowing its history 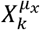. [2] From Eq. 1, we get that:

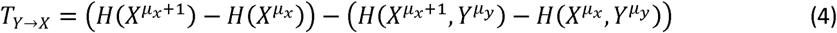

where *H* denotes the Shannon entropy of either process 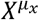 or 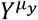, or the joint processes 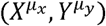 or 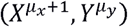, defined as [13, 14]

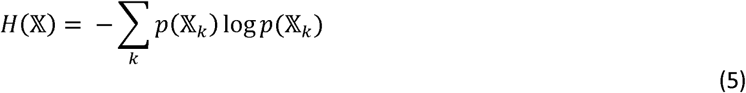

where the sum runs over all states.

Eq. (4) shows that T_*Y*→*x*_ equals the conditional mutual information, 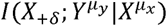 [15]

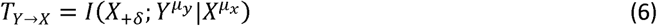

Eq. (6) indicates that the transfer entropy, T_Y→x_ represents the average amount of information contained in the source about the next state X_+ δ_ of the destination, that was not already contained in the destination’s past.

### Local Transfer Entropy

To determine a local transfer entropy measure, we first note that Eq. (3) is summed over all possible state transition tuples 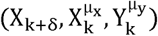, weighted by the probability of observing each of such tuple, 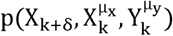. Therefore, we can write: [16]

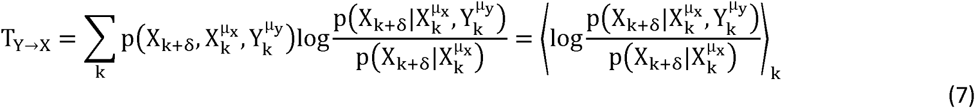

where ⟨ … ⟩_*k*_ indicates the average with probability 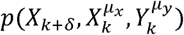 overall states *k* of the following quantity:

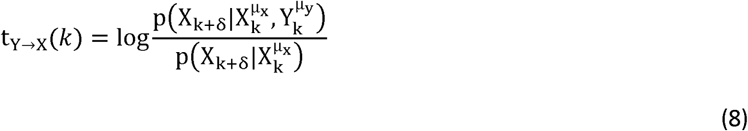

Here, the quantity *t*_*Y*→*x*_ (*k*) denotes the so-called *local transfer entropy* at the time step *k* [16] Eq. (8) can be further simplified as:

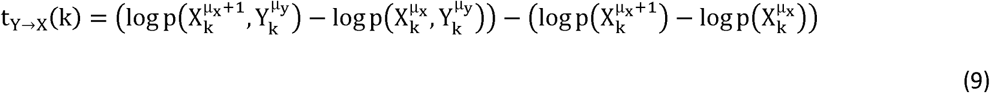

### Mutual Information

Based on the information theory [13], the mutual information, *I(X;Y)*, equals the so-called *Kullback-Leibler distance,K*_*D*_ [17]:

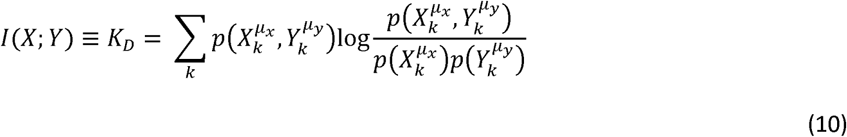

*I(X;Y)* is a measure of the dependence between two random variables [13]. Eq. (10) can also be expressed in terms of the Shannon information entropy as the following [13]:

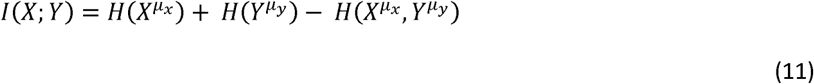

### Effective Transfer Entropy

The probability distributions of the time series are not known a *priory*, and hence, the computation of the transfer entropy is complicated and subject to noise [15]. Besides, for large time lags *τ*_*x*_ and *τ*_*y*_, a fairly wide joint distribution will be obtained. Because of this, the transfer entropy can be different from zero, even when there is no causal relationship between two random processes (see, for example, Ref. [2] and the references therein.) To avoid that the so-called *effective transfer entropy* 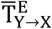 is computed based on data shuffling [15] [18], which is defined as [2]:

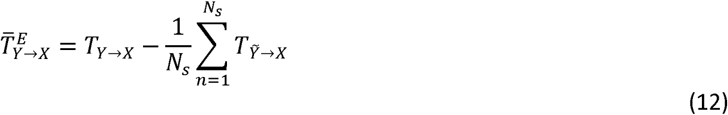

where 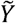 represents the shuffled time series and *N*_*s*_ denotes the number of random shufflings. The main aim is that by shuffling the time series *X*, or *Y*, we entirely removes any dependence between them due to biasing, without changing their distributions. [2] This means that the transfer entropy, calculated from the shuffled time series, can serve as a significance threshold to distinguish information flow. That is, we can exclude any dependency on the information transfer between two coupled systems introduced by external factors, such as added bias or noise. Therefore, the following term is a considered as a measure of the bias term in the transfer entropy:

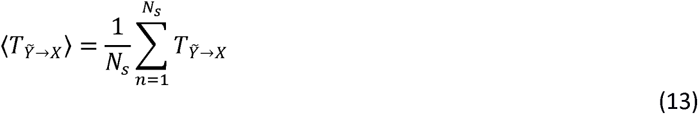

Thus, if there is no added bias or noise in the dynamics of the real processes, then there is no causal relationship between *X* and 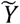, and hence 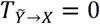, otherwise 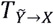 will capture any causal relationship due to the added bias or noise in the real dynamics of variable at different time steps. Therefore, Eq. (12) can also be interpreted as the following relationship:

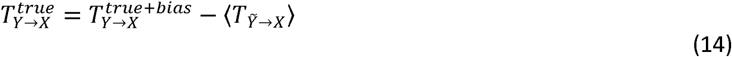

where 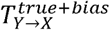 is a single attempt of computing transfer entropy from two input time series representing, for example, two components of a dynamical system, which may incorporate bias or noise in their dynamics and 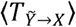 is computed using Eq. (13), which attempts to remove the added bias or noise in the system. Then, the hope is that the difference between these two terms will be a measure of the true causal relationship between the two processes, 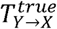, originating from the real dynamics.

That has the advantage that the new measure of the transfer entropy, introduced previously [2] (and the references therein), can capture system dynamics that traditional measures cannot.

## 3 Algorithm Design using Symbolic Analysis

### Analyzing Collective Variables Using Machine Learning Approach

The dynamical variables characterizing either different components of a system or different dynamical systems describe the collective or essential degrees of freedom of the system. These collective variables are often determined using the principal components analysis (PCA) [19] as for biomolecular systems [20]. We have discussed in details the PCA algorithm in Ref. [21] In this study, we aim to introduce a new algorithm, which is an improved version of the auto-encoder machine learning approach [22] to the algorithm of determining the collective variables from higher dimensionality data. Machine Learning (ML) approach provides a potential method to predict the properties of a system using decision-making algorithms, based on some predefined features characterizing these properties of the system. There exist different ML methods used to predict missing data and discover new patterns during the data mining process. [23] Artificial neural network (ANN) methods consider a large training dataset, and then it tries to construct a system, which is made up of rules for recognizing the patterns within the training data set by a machine learning process. [24]

In general, for an ML process based on the ANN contains *K* hidden layers (see also Figure 1), the output *Z*_*i*_ of each layer is defined in terms of the input *X*_*i*_ as

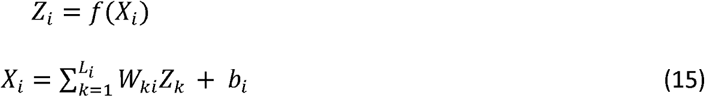

**Figure 1:**
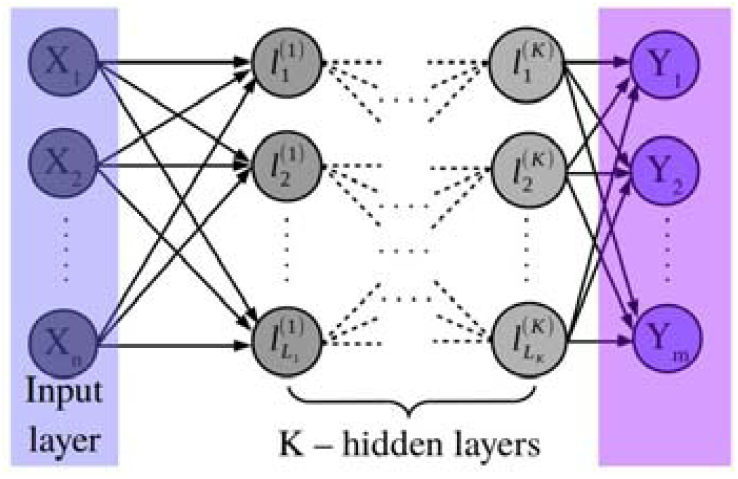
Illustration diagram of an ML process based on the ANN, characterized by an input vector of dimension *n, K* hidden layers of 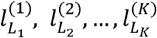 neurons each, and an output vector of dimension m. [24]

Here, *f* is often taken to be a sigmoid function, *W*_*ki*_ is the weight matrix determining the connectivity between the neurons and *b*_*i*_ is the bias term.

Here, ***W*** and ***b*** are free parameters, which need to be optimized for a given training data used as inputs and given outputs, which are known. To optimize those parameters, the so-called loss function is minimized using the Gradient Descent method [24]:

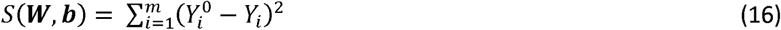

***Y***^0^ represents the true output vector. For that, the derivatives of *S*(***W***,***b***) for ***W*** and ***b*** are calculated:

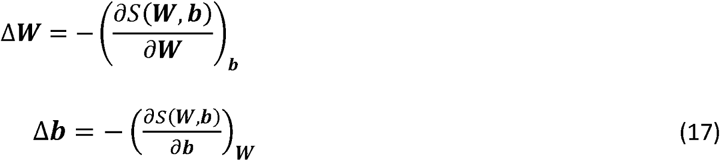

The following regularization terms avoid overfitting, which is one of the pitfalls of the machine learning approaches [25]:

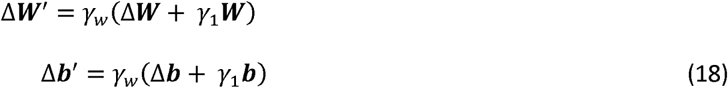

Here, *γ*_*w*_ is called learning rate for the gradient and *γ*_1_ is called the regulation strength. Because the Gradient Descent method often converges to a local minimum, it provides a local optimization to the problem. To avoid this pitfall, we are going to introduce a new approach, called here as Bootstrapping Swarm Artificial Neural Network (BSANN). [24]

The standard ANN method deals with random numbers, which are used to initialize the parameters ***W*** and ***b***; therefore, the optimal solution of the problem will be different for different runs. In particular, we can say that there exists an uncertainty in the calculation of the optimal solution (i.e., in determining ***W*** and ***b***.) To calculate these uncertainties in the estimation of the optimal parameters, ***W*** and ***b***, we introduce a new approach, namely bootstrapping artificial neural network based on the method proposed in [26]. In this approach, M interacting copies of the same neural network are run independently using different input vectors. Then, at regular intervals, we swap optimal parameters (i.e., ***W*** and ***b***) between the two neighboring neural networks.

Furthermore, to achieve a good sampling of the phase space extended by the vectors ***W*** and ***b***, we introduce two other regularization terms similar to the swarm-particle sampling approach. First, we define two vectors for each neural network, namely 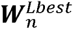 and 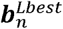, which represent the best local optimal parameters for each neural network. In addition, we also define ***W***^*Gbest*^ and ***b***^*Gbest*^, which represent the global best optimal parameters among all neural networks.

Then, the expressions of the gradients modify by introducing two additional regularization terms for each neural network configuration *n, n* = 1,2, …, *M* like the following: [24]

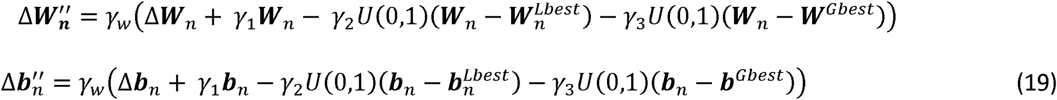

Here,*U* (0,1) is a random number between zero and one, and *γ*_2_ and *γ*_3_ represent the strength of biases toward the local best optimal parameters and global best optimal parameters, respectively. The first term indicates the individual knowledge of each neural network and the second bias term the social knowledge among the neural networks. Then, the weights, ***W***_*n*_, and biases, ***b***_*n*_, for each neural network *n* are updated at each iteration step according to:

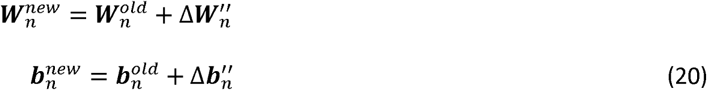

Consider the vector ***Q***^*T*^, which represents *T* time frames of a dynamical system or one of its components:

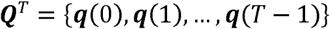

where ***q***(*t*) (for *t* = 0,1, …, *T* − 1) represents a configuration of the system (or its components) of *g* degrees of freedom:

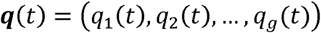

That forms a Markovian chain of the states of a stationary stochastic random process visited by the dynamical system. The problem is to find a reduced *g*′ dimensional space (*g*′ < *g*), which compresses the data. This is suggested here by determining an encoding function as

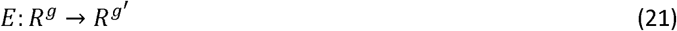

and a decoding function as the following:

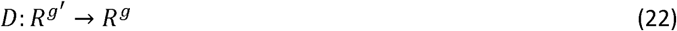

The function *E* provides a non-linear mapping using the Bootstrapping Swarm Artificial Neural Network of the Cartesian coordinates ***q***(*t*) as:

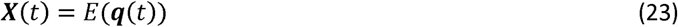

where ***X*** (*t*) is an *g*′-dimensional vector in the essential subspace of slow collective variables representing either the entire system or its components. Then, similarly, using the non-linear mapping *D* we obtain an approximate time-lagged signal, 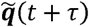:

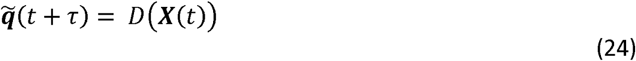

That aims at average to minimize the error using the variation principle:

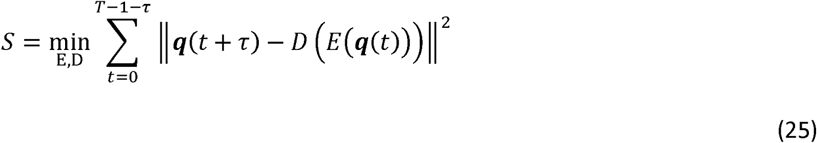

Here, *τ* is the time-lag of the input signal ***q***(*t*), and the approach is called time-lagged auto-encoder. For *τ* = 0, the approach represents the standard auto-encoder method. Note that both the input and output signal of the encoder-decoder non-linear neural network is the trajectory ***q***(*t*) in the Cartesian space and the output signal of the encoder, which is the input signal for the decoder, represent the slow collective variables ***X***(*t*).

To create the input signals (see [22] for the details of the algorithm), first, two new signals are reconstructed using the Cartesian space vectors:

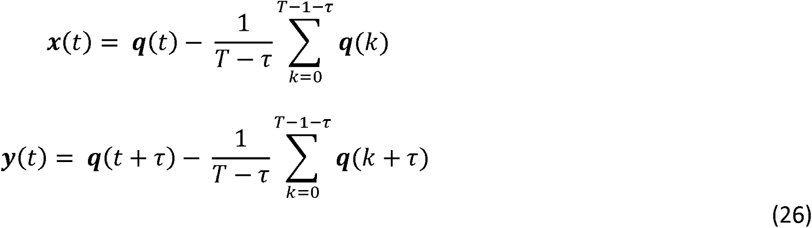

The covariance matrices are constructed as the following:

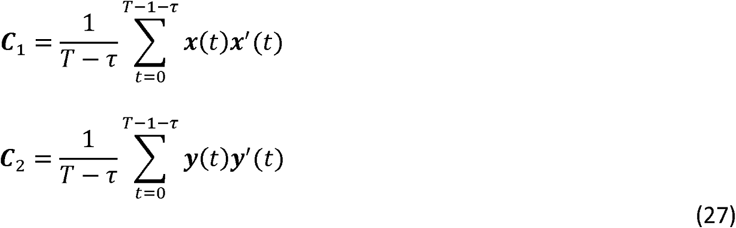

where (′) denotes the transpose of a vector. Then, both signals ***x***(*t*) and ***y***(*t*) are whitened as the following:

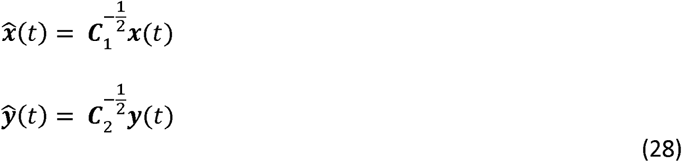

These two signals are the input and the output, respectively, of the encoder-decoder approach, which aims to define the non-linear functions *E* and *D* (that represent the BSANN algorithm), such that, the following reconstructed error is minimum:

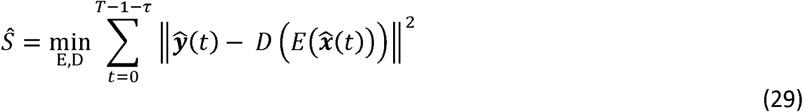

Here, the input signals ***q***(*t*) could characterize the dynamics of the entire dynamical systems, such as in the applications of measurements of the information flow in signal pathways. Besides, ***q***(*t*) could characterize the dynamics of a component of the dynamical system, such as in the applications of finding the information flow between different components of the same system; for example, it may represent the atomic coordinates of each amino acid in a protein structure in determining the information flow between different amino acids of a protein.

### Symbolic Analysis

In this study, the symbolic analysis approach is followed based on coarse-graining of the time series into symbols [27, 28, 16, 2, 12]. In particular, we use the symbolization technique proposed in Ref. [2], which we found to be computationally very robust and at the same time, maximizing the information content about the real-time series. This approach creates the following symbolic sequence for each time series (*X*_0_,*X*_1_, …,*X*_*N*−1_):

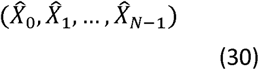

This process is also called *coarse-graining* [2] (and the references therein). In the coarse-graining, all information concerning the dynamics of series is suitably encoded using partitioning of phase space.

The time-series (*X*_0_,*X*_1_, …,*X*_*N*−1_) is converted into a symbolic sequence using the following rule [2]

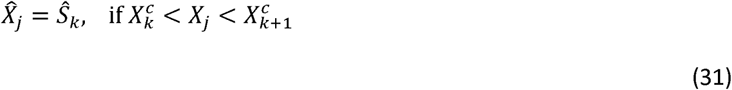

where 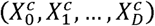 is a given set of *D*+1 critical points, and (*Ŝ*_0_, *Ŝ*_1_, …, *Ŝ*_D−1_) is a set of *D* symbols, here the numbers 0,1,2, … *D* − 1. Here, *D* is such that the Kraft inequality is satisfied [13]:

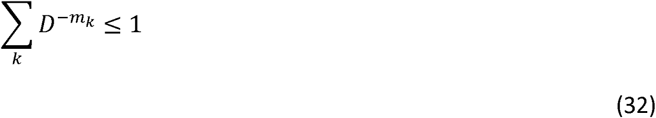

where the sum runs over all state vectors, and *m*_*k*_ is the length of each state vector, which for a single time series is considered to be equal for every state vector. The new state vector generated in this way,

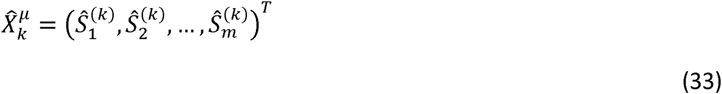

represents a symbolic state vector, which is a subset of numbers from 0 to *D* − 1. Concatenation of the symbols of a sequence of length m yields the word *W*_*k*_:

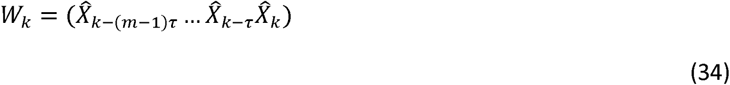

A particular sequence of symbols 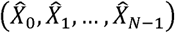 is uniquely characterized by the words *W*_*k*_ for *K* = *K*_0_, …,*N* − 1. [2] The probability of finding a particular value of *W*_*k*_, is calculated from the simulation data, and used to compute the Shannon entropy,

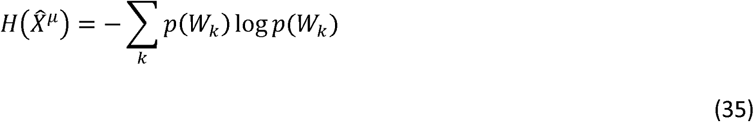

where the sum runs over all symbolic state vectors represented by the words *W*_*k*_. Since the time series (*X*_0_,*X*_1_, …,*X*_*N*−1_) is mapped onto the symbolic sequence 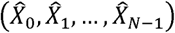 uniquely (i.e., the symbolic representation is injective) the entropies 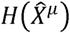 and *H*(*X* ^*µ*^) coincides [29].

We obtain the critical points 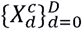 for a particular series by maximizing the entropy for all possible partitions. [2] Increasing the number of critical points will initially increase the information entropy, but after a sufficient number of critical points, the information entropy plateaus. At this point, the optimum number of critical points has been reached; a further increase will not increase the accuracy of the calculation but does slow down the computation. [13]

In our implementation, we optimize critical by maximizing the Shannon entropy through a Monte Carlo approach [2]. Similarly, in this study, it has been proved that the joint Shannon information entropy of two discrete symbolic processes 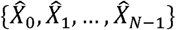 and {Ŷ_0_, Ŷ_1_, …, Ŷ_N−1_}can be calculated as,

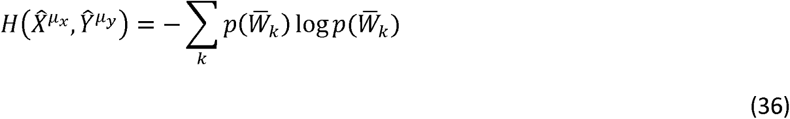

where 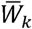 is the concatenation of two words, 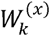 and 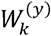, representing the words of processes *X* and *Y*, respectively.

In general, the length of discrete processes and the number of states are limited by the sampling. To correct for the finite sampling of the time-discrete processes, we use [11] as implemented in Ref. [2]:

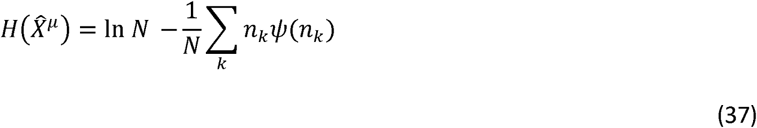

Where the sum is over all states,*n*_*k*_ is the frequency of observing state *k*, and *ψ* (*x*) is the derivative of Gamma function *Γ* for *x*. Using the above-described coarse-graining of the time series approach, the *symbolic transfer entropy* can be written as [2]:

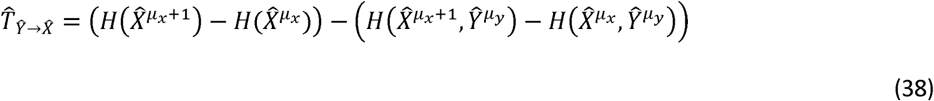

Similarly, in this study, we have defined the so-called *symbolic local transfer entropy* as:

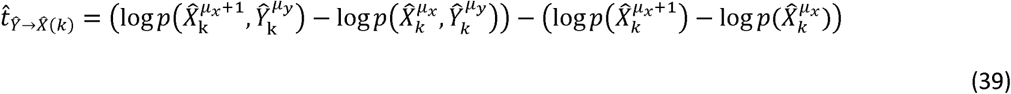

In addition, *symbolic mutual information Î*(*X; Y*) is calculated as:

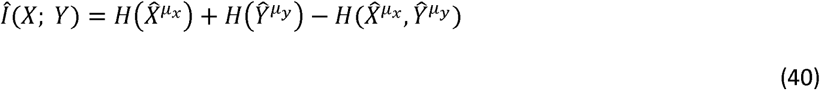

### Algorithm for Optimization of Embedded Parameters

With the proper values of *m* and *τ* embedding parameters, a smooth discrete-time process is defined, which reconstructs the underlying dynamics. [2] The choice of these two parameters is crucial for the proper characterization of the structure of the time series [30, 31, 32]. The mathematical concepts for defining the state vector dimension *m* have been reviewed in details in Refs. [33, 12]. Several methods have been proposed for estimating the optimal embedding parameters *m* and *τ* simultaneously [31], which are based on minimizing the number of false nearest neighbors. A comparison of these methods has been reported [34], as well as an approach that combines the global false nearest neighbor (GFNN) method for the calculation of *m*, with a separate method for determining the time lag, *τ*, using the autocorrelation function to determine *τ* as its first zero. Here, for estimation of the embedding parameters *m* and *τ* we implemented the algorithm presented in Ref. [2]. In particular, the time lag *τ* is determined as the first minimum of the mutual information written as a function of the time lag [2] (and the references therein):

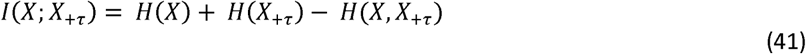

On the other hand, space dimension *m* is determined from a separate procedure using the false nearest neighbors. For that, given a state vector by Eq. (1), the nearest neighbor is

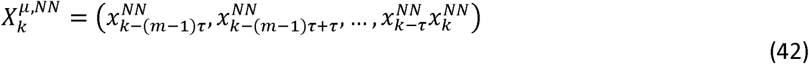

The Euclidean distance, between these two points in m-dimensional space, is given by

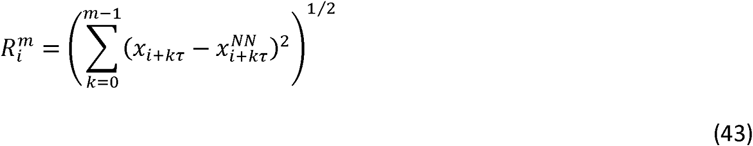

The distance between these two points in the (*m*+1) –dimensional space is,

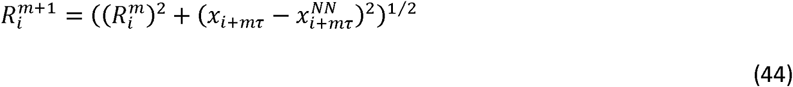

This distance is normalized against the distance in *m*-dimensional space:

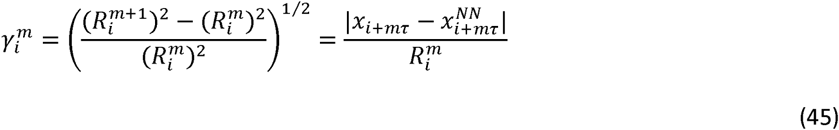

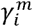 is compared to a threshold value *R*_*tol*_, which is determined a priori and recommended to be 15 [34]. In this study, we tested the algorithm for different values of *R*_*tol*_ in the range [2].

If 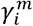 exceeds *R*_*tol*_, then 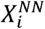 is the false nearest neighbor of *X*_*i*_ in the *m*-dimensional space and *f*_*FNN*_, the frequency of the false nearest neighbors is increased by one. The value of *m* is increased until *f*_*NN*_ approaches zero. [2]

## 4 Software Framework

### 4.1 Software Architecture

A flowchart diagram of the software is shown in Figure 2. It contains five main modules, namely EMBD_CLASS, SYMB_CLASS, TE_CLASS, LTE_CLASS, and MI_CLASS, which are described in the following.

**Figure 2:**
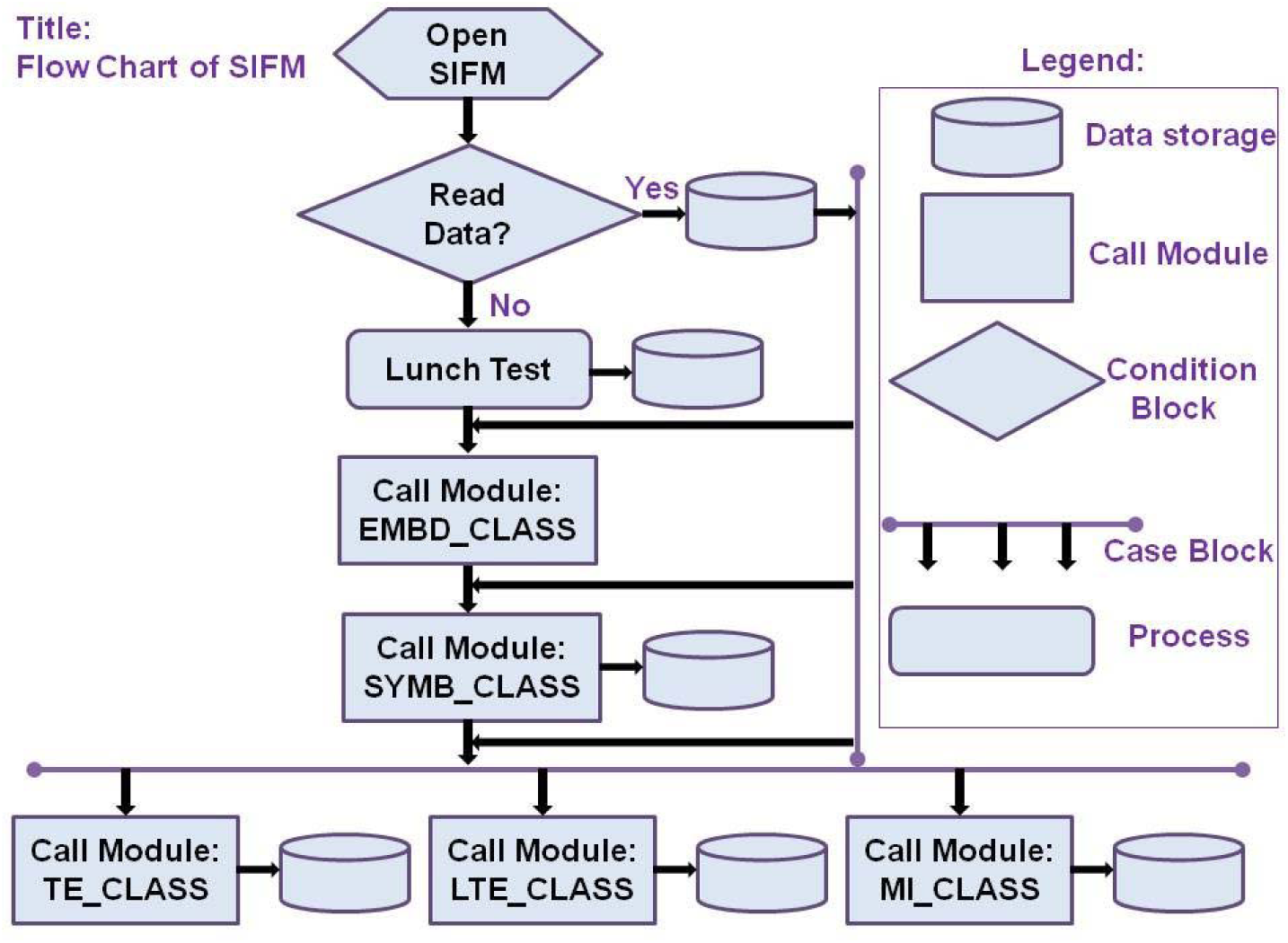
A flowchart diagram of the SIFM software.

#### Embedded Parameters Module (*EMBD_CLASS*)

This class contains the functions and subroutines for computation of the embedded parameters,m and τ. A UML diagram of this module is shown in Figure 3. As can be seen, this class inherits the methods from three auxiliary modules, respectively, SIFM_CLASS, TEUTILS_CLASS, and TERANDOM_CLASS, which are described in the following.

**Figure 3:**
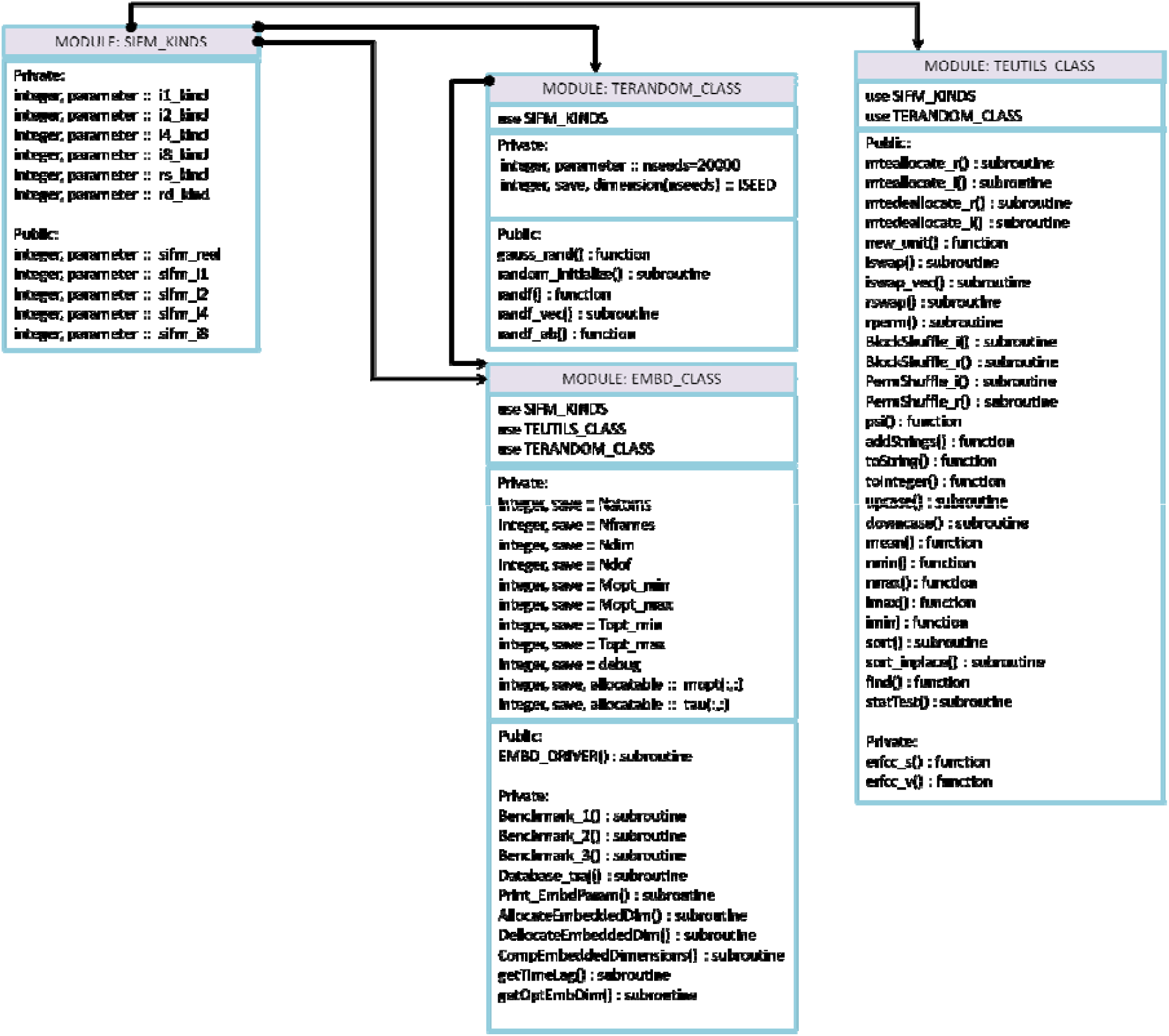
UML diagram of the module EMBD_CLASS.

#### Symbolize Trajectory Module (*SYMB_CLASS*)

This class contains the subroutines and the functions used for creating of the symbolic trajectory using different methods. Figure 4 presents a UML diagram of this class along with auxiliary modules inherited by the SYMB_CLASS.

**Figure 4:**
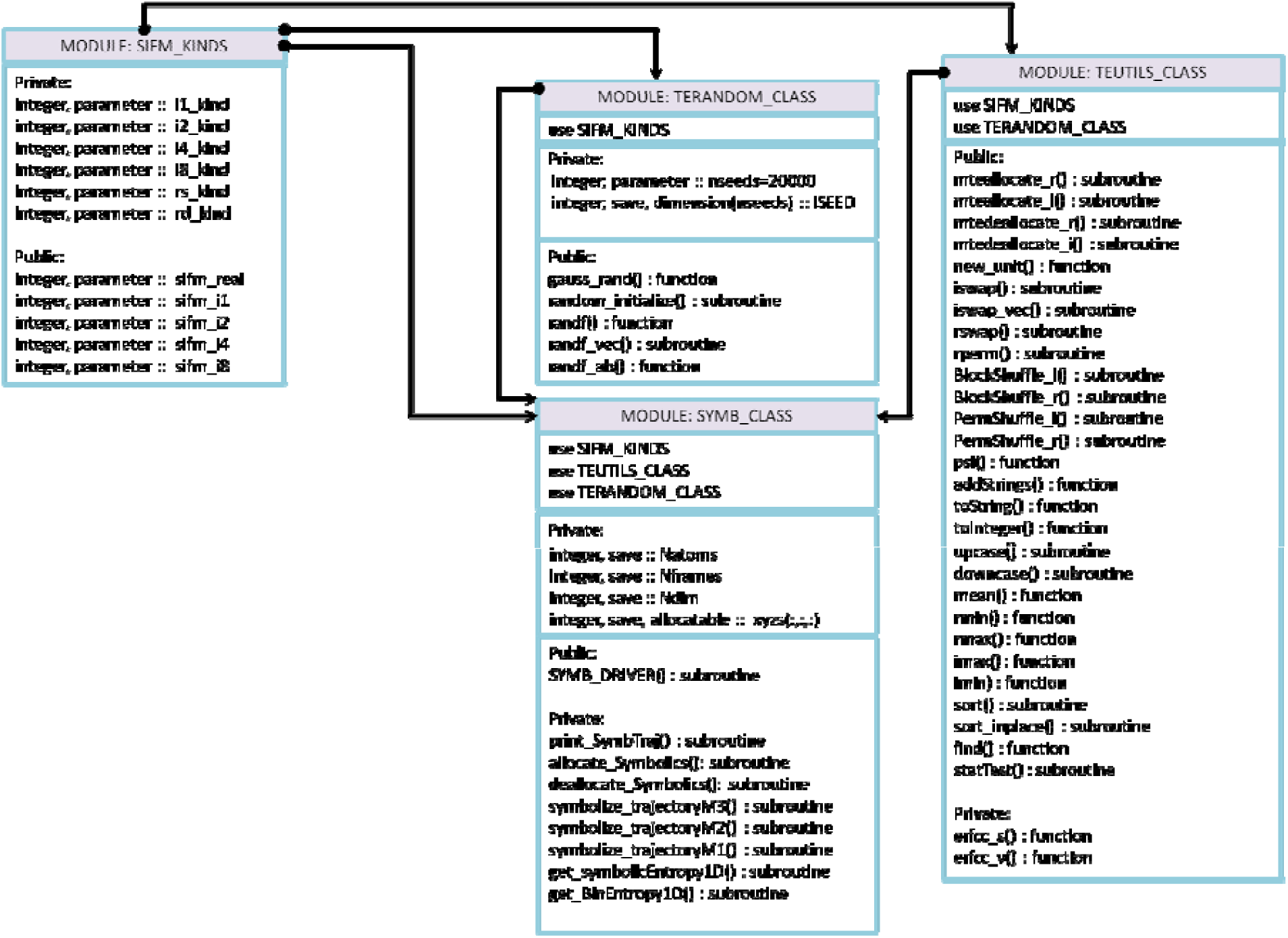
UML diagram of the module SYMB_CLASS.

#### Symbolic Transfer Entropy Module (*TE_CLASS*)

This class contains the subroutines and the functions used for calculations of the symbolic transfer entropy. In Figure 5, we have presented the UML diagram of TE_CLASS module. Besides, we have shown the auxiliary modules, including TELINKLIST_CLASS, which will be described in the following.

**Figure 5:**
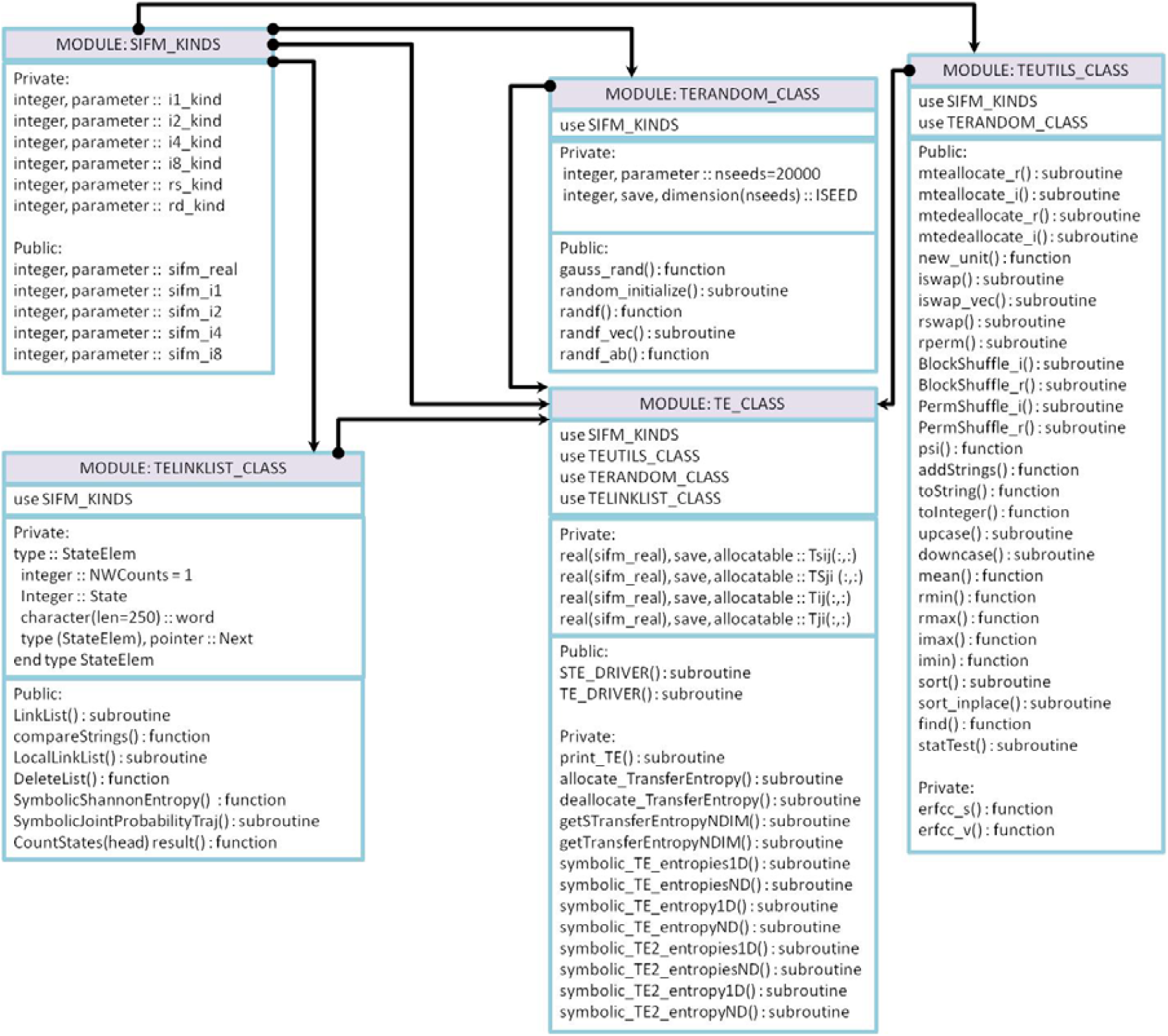
UML diagram of the module TE_CLASS.

#### Symbolic Local Transfer Entropy Module (*LTE_CLASS*)

This class contains the subroutines and the functions used for calculations of the symbolic local transfer entropy. In Figure 6, we have presented the UML diagram of LTE_CLASS module. In addition, we have shown the auxiliary modules called by LTE_CLASS.

**Figure 6:**
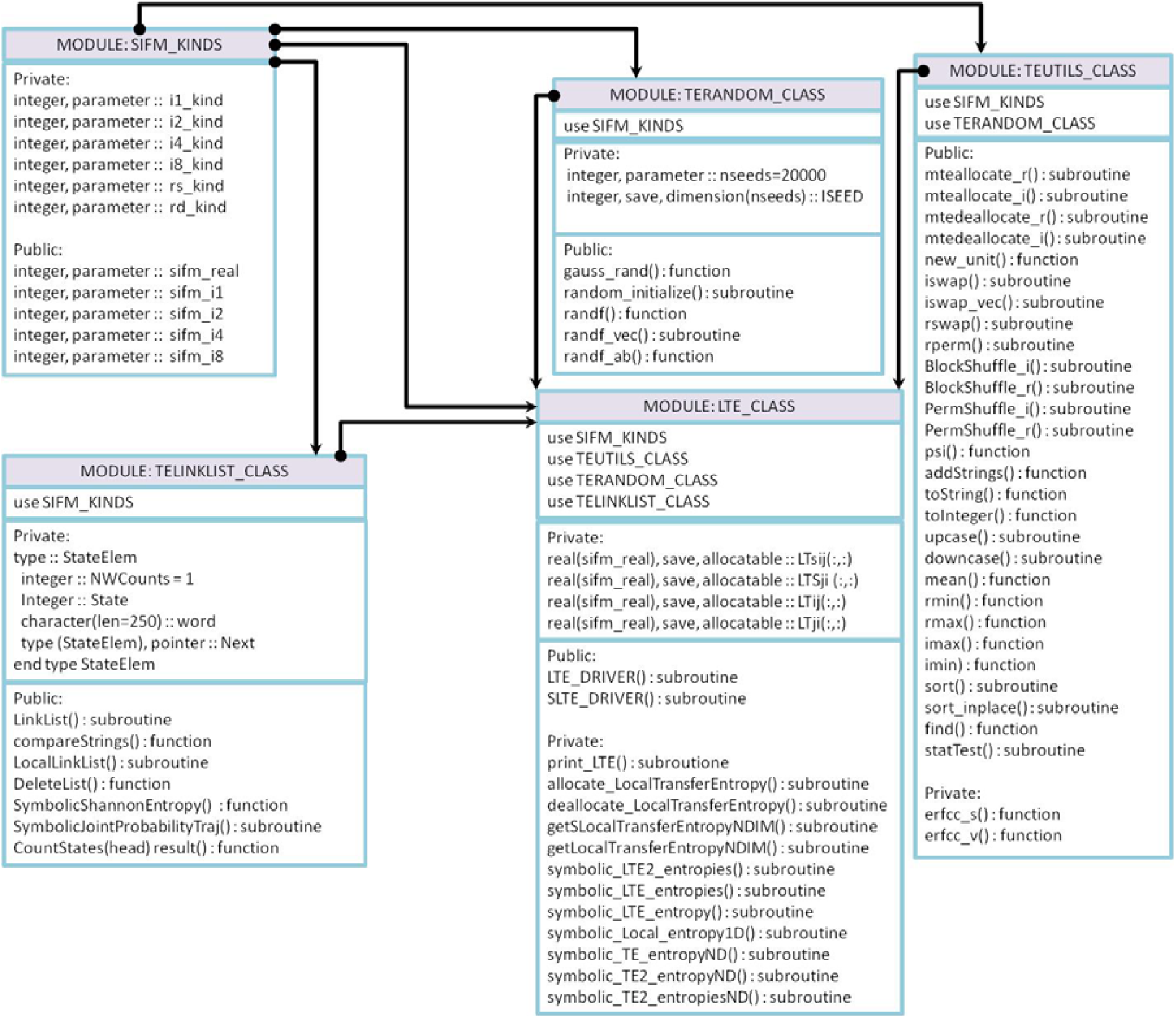
UML diagram of the module LTE_CLASS.

#### Symbolic Mutual Information Module (*MI_CLASS*)

This class contains the subroutines and the functions used for calculations of the symbolic local transfer entropy. Figure 7 shows the UML diagram of the MI_CLASS module. Also, we have shown the auxiliary modules called by MI_CLASS.

**Figure 7:**
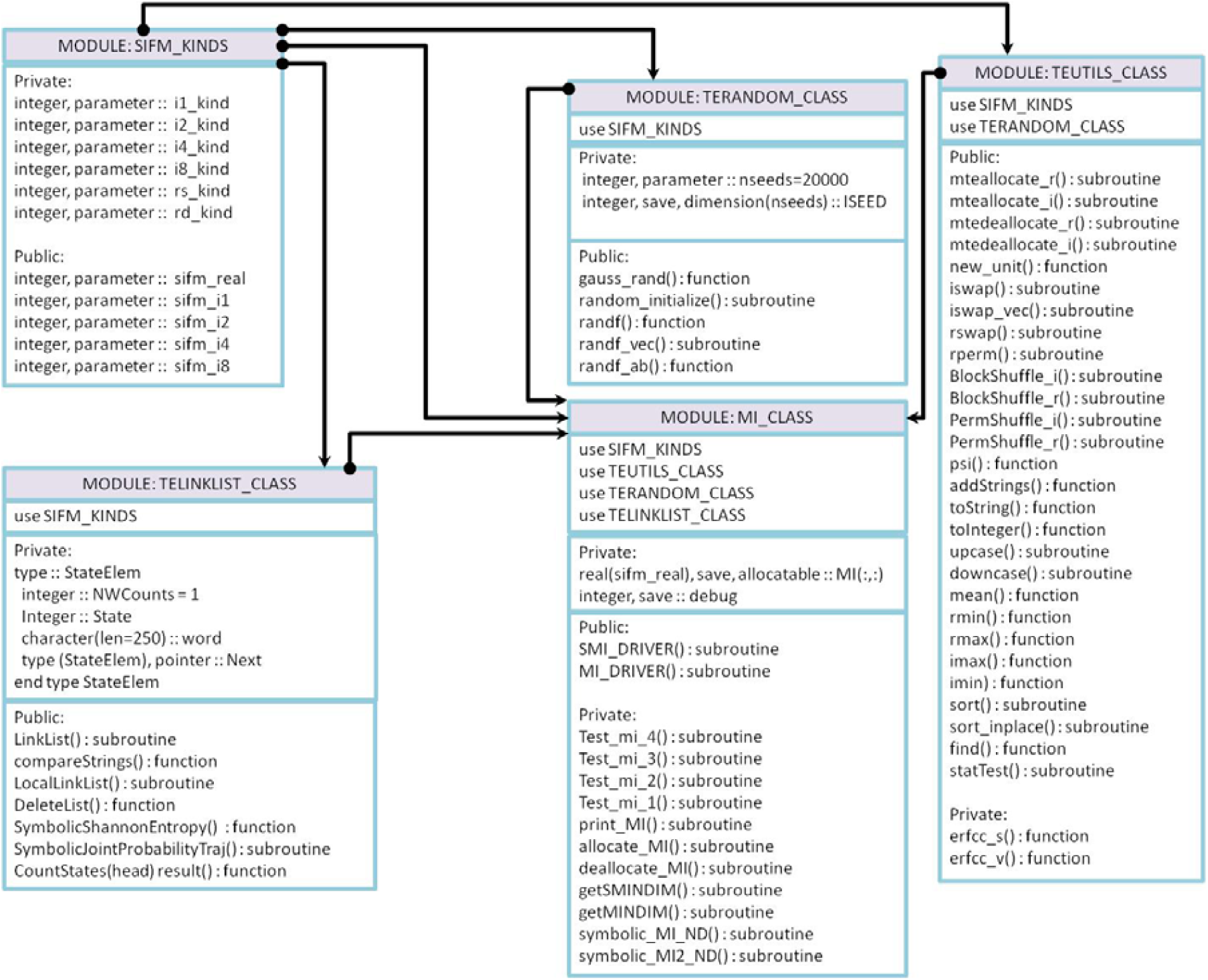
UML diagram of the module MI_CLASS.

#### Auxiliary Class Modules

There are four auxiliary class modules, namely SIFM_KINDS, TEUTILS_CLASS, TERANDOM_CLASS, and TE_LINKLIST_CLASS modules. Figure 8 presents the UML diagram of the auxiliary modules, SIFM_KINDS, TEUTILS_CLASS, TERANDOM_CLASS, and TE_LINKLIST_CLASS and the relationship between them as depicted by arrows.

**Figure 8:**
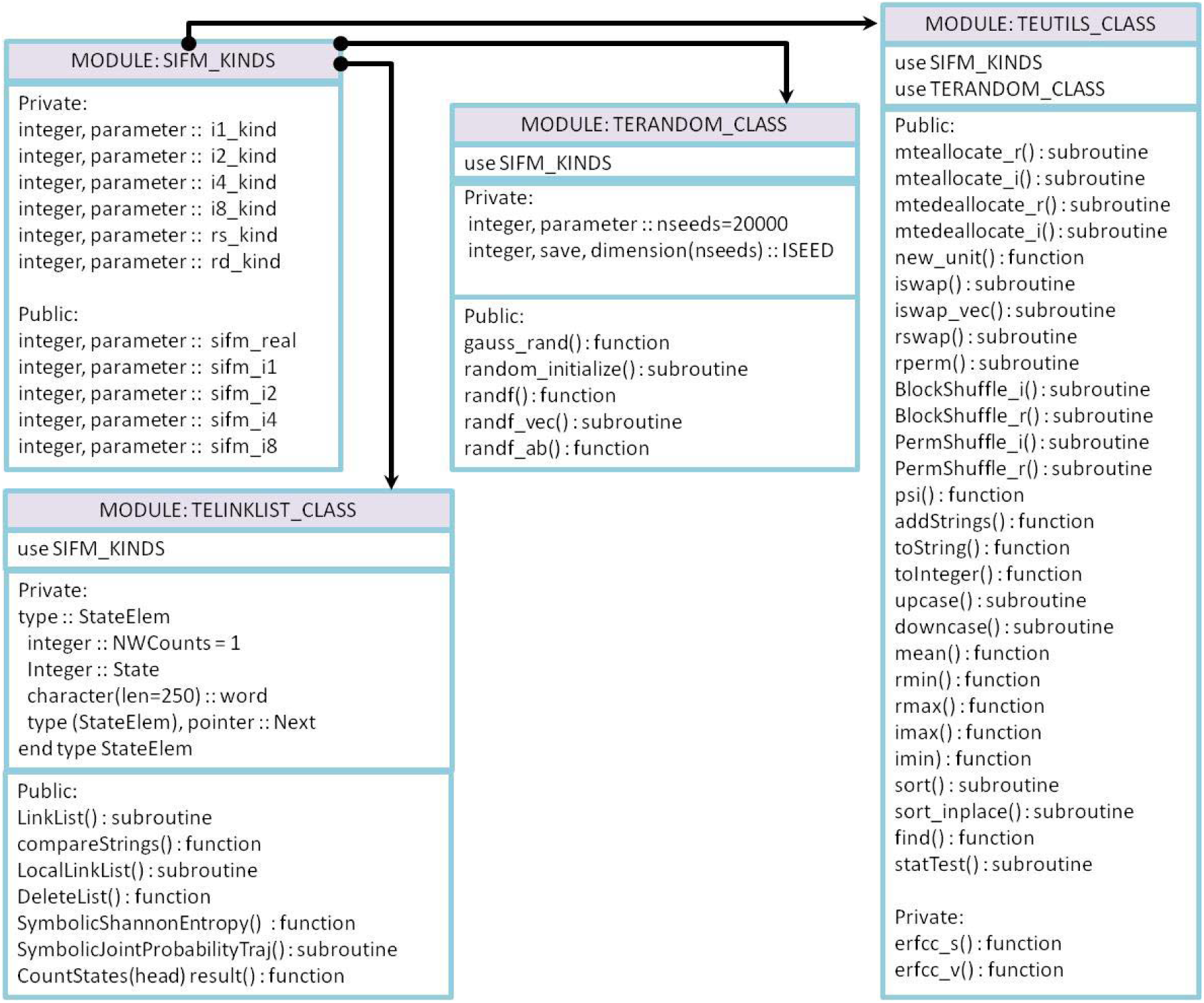
UML diagram of the auxiliary modules.

### 4.2 Software Functionalities and Implementation

#### Portability

SIFM is written in Fortran 90 object-oriented style, making it very portable in (almost) all operating systems.

#### Computational Precision

The precision of the computation can be adjusted by the user using KIND variables as described in the module SIFM_KINDS, for example, by adjusting the variables *sifm_real, sifm_i1, sifm_i2, sifm_i4, or sifm_i8*. Besides, SIFM uses fast intrinsic routines of Fortran for manipulating the operations with strings or converting integer symbols into string symbols.

#### Memory Management

The software is written in the modular form, making it faster to compile and better memory management. Moreover, all the dynamical arrays are declared as either allocatable arrays or pointers, which provide efficient management of the program memory.

#### Data Management

In SIFM, each module produces its data output, which is used either for further analysis or by other modules as input information. That is in particular important when the program is interrupted or crashed due to errors, and then a new restart can be set up from the last point. Also, SIFM supports different kind of input data format, including binary files, for reducing the size of the data files. Furthermore, SIFM supports the majority of the trajectory file formats that are produced by other software as well, making this software applicable to many other fields of applications.

#### Parallelization

SIFM is highly parallelized using Message Passing Protocols (MPI). In this version of the software, we have parallelized the computation of symbolic trajectories using the Monte Carlo technique and the computation of symbolic transfer entropy, local transfer entropy, and mutual information. That is in particularly useful when performing symbolic analysis of more than one pair of time series, let say of *n* pairs,

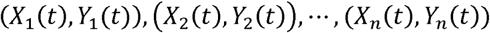

Then, since the computations within pairs are independent, we can distribute these computations among processors, let say, *P* processors, as illustrated in Figure 9. Here, every processor is performing *m* calculations with *m*= *P*/*n.* In order to guarantee an equal load balance between the processors, *P* should be chosen such that *P*/*n* is an integer number.

**Figure 9:**
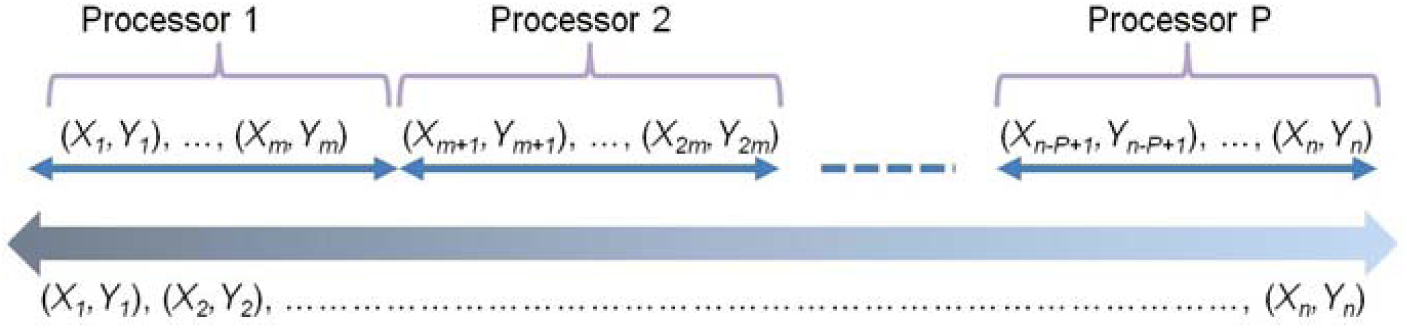
Illustration of the workload distribution for the symbolic computation of *n* pairs of time series among the *P* processors.

In Figure 10, we show the average CPU time (in seconds) for computation of embedded parameters(m, τ) per degree of freedom (i.e., time series) as a function of the number of available processors and length of the time series. The averages are carried out over different dynamical systems. It shows that CPU time decreases proportionally with the number of processors, and as expected, it increases with the length of the time series. Typically, it takes 2.3 seconds to compute both m and τ for a time series of length 20000 frames using 8 CPUs.

**Figure 10:**
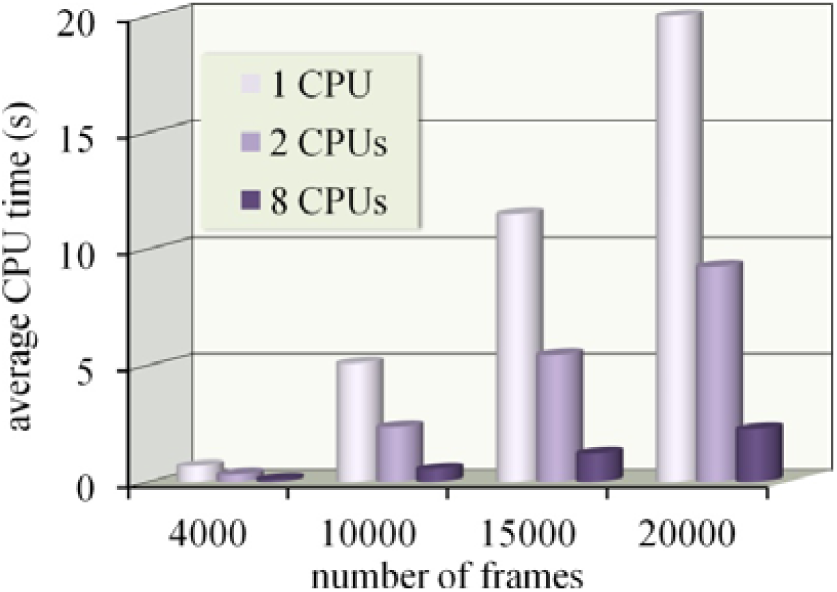
Average CPU time (in seconds) for the computation of embedded parameters (*m, τ*) per time series as a function of the length of the time series and number of processors. The averages are carried out over different time series.

In Figure 11, we show the speedup of the parallel computations for symbolic transfer entropy and symbolic mutual information versus the number of processors. For the sake of comparisons, the ideal expected speed up is also shown. It shows that up to four processors, the estimated speedup is the same as the ideal one. Deviations from this linearity are observed with the further increase of the number of processors due to overhead time on the master node. Note that these computations are performed on the i7 computer architecture.

**Figure 11:**
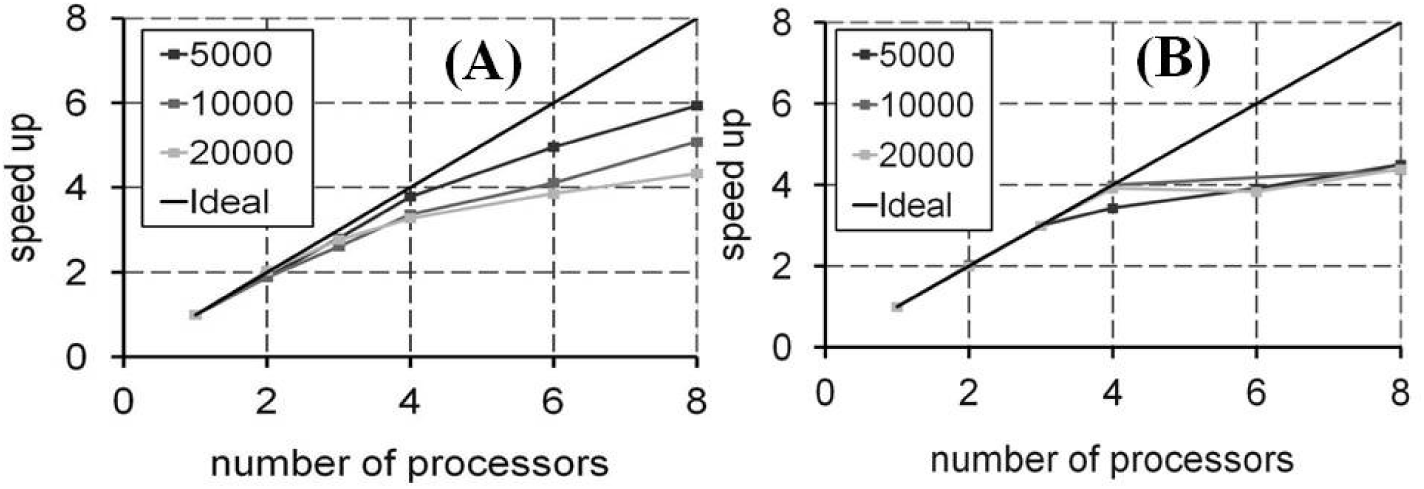
Speed up of (A) Symbolic Transfer Entropy and (B) Symbolic Mutual Information calculations as a function of the number of processors for the different lengths of time series.

#### Graphical User Interface

SIFM has a Graphical User Interface (GUI) implemented using Matlab. For example, in Figure 12, we have shown the GUI for calculation of the embedded parameters using EMBD_CLASS module. Four parameters are needed to compute these parameters, namely the minimum and maximum values of the time lag,, and embedded dimension,. In addition, the GUI has the functionality of plotting the results as graphics, which then can be exported in a separate figure and saved for later use (see Figure 12).

**Figure 12:**
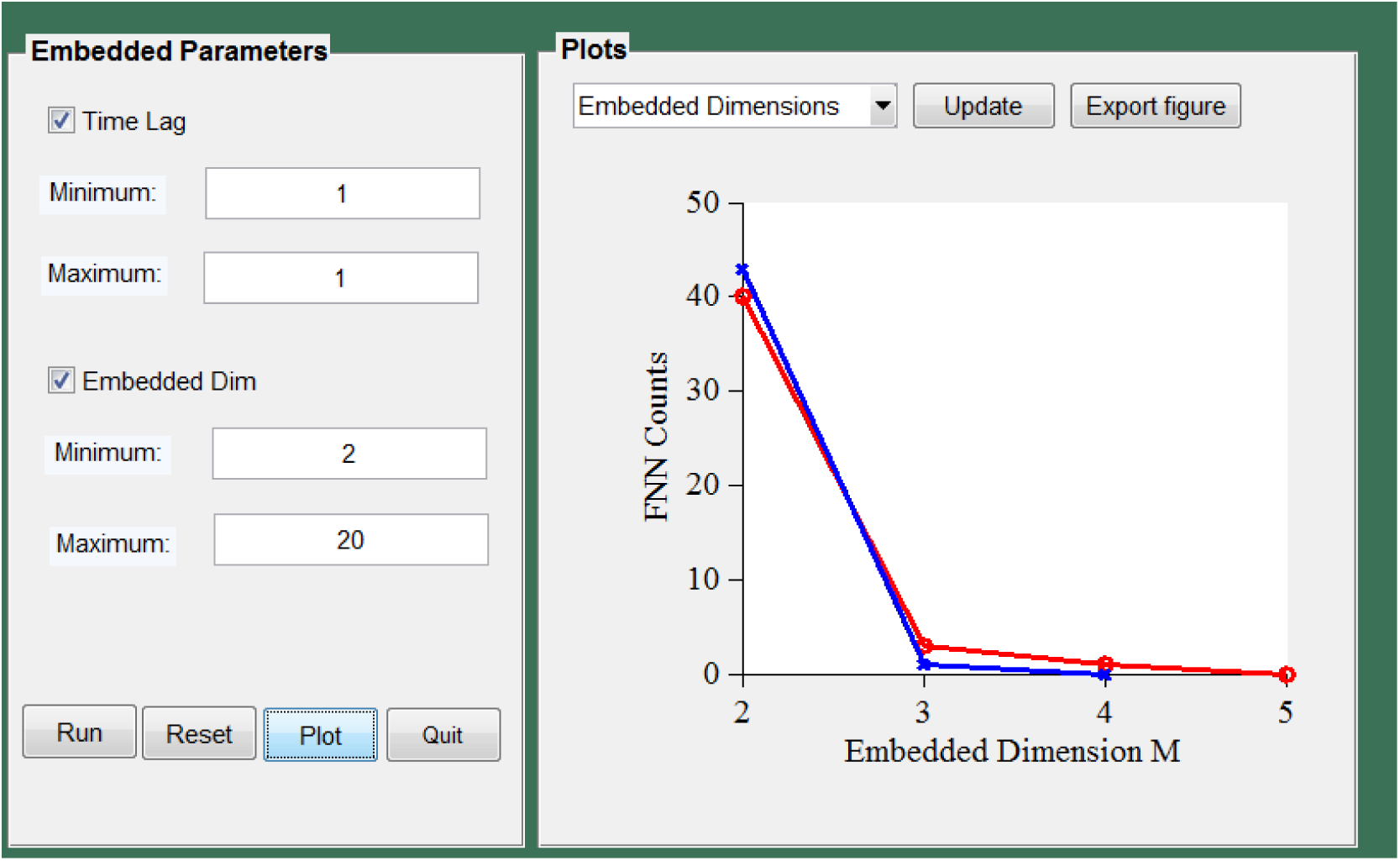
GUI for calculation of the embedded parameters using EMBD_CLASS module.

## 5 Illustrative Examples

### Benchmark 1

The coupled noisy time series is as the following:

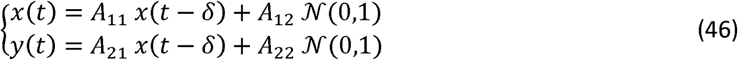

where *A*_11_, *A*_12_, *A*_21_, and *A*_22_ are constants and *𝒩* (0,1) is a random number distributed according to a normal distribution with mean zero and variance one. In Eq. (31), the coefficient *A*_21_ is the strength of coupling between the random variables *X* and *Y*, and *A*_22_ the strength of the external noise on variable *Y. δ* indicates a time shift in the history dependence, which is taken here to be one. In Figure 13, we show the symbolic transfer entropies as a function of the coupling strength. *A*_21_ The values of other parameters are fixed, as shown in Figure 13.

**Figure 13:**
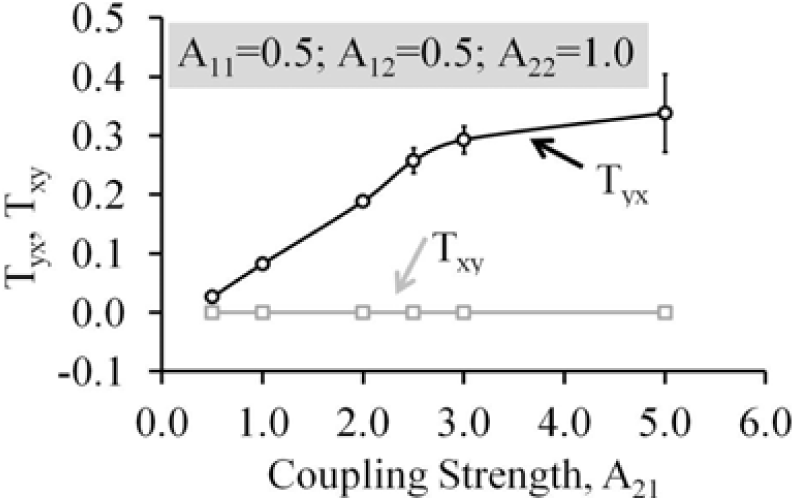
Symbolic transfer entropies, *T*_*xy*_ and *T*_*yx*_ versus the coupling strength *A*_21_ for fixed choices of other parameters as depicted in the figure.

### Benchmark 2

This benchmark corresponds to the C2-Fc complex biomolecular system, where C2 is a fragment of protein G, and Fc is a domain of human IgG protein.C2 fragment has 56 amino acids, and Fc has 206 amino acids. We performed molecular dynamics simulations of the complex for 30 ns. The first ten ns are omitted from the analysis, and only the last 20 ns are used for calculations. We printed out the configurations every two ps, thus, in total, 10000 snapshots were used for calculations of the symbolic transfer entropies. In Figure 14, we show the directional symbolic transfer entropies, *D*_i→j_, as a color map for the C2 fragment (Figure 14A), Fc fragment (Figure 14B) and between C2 and Fc (Figure 14C). Here, the directional symbolic transfer entropy between two-time series *X* and *Y* is calculated as

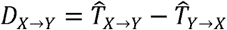

**Figure 14:**
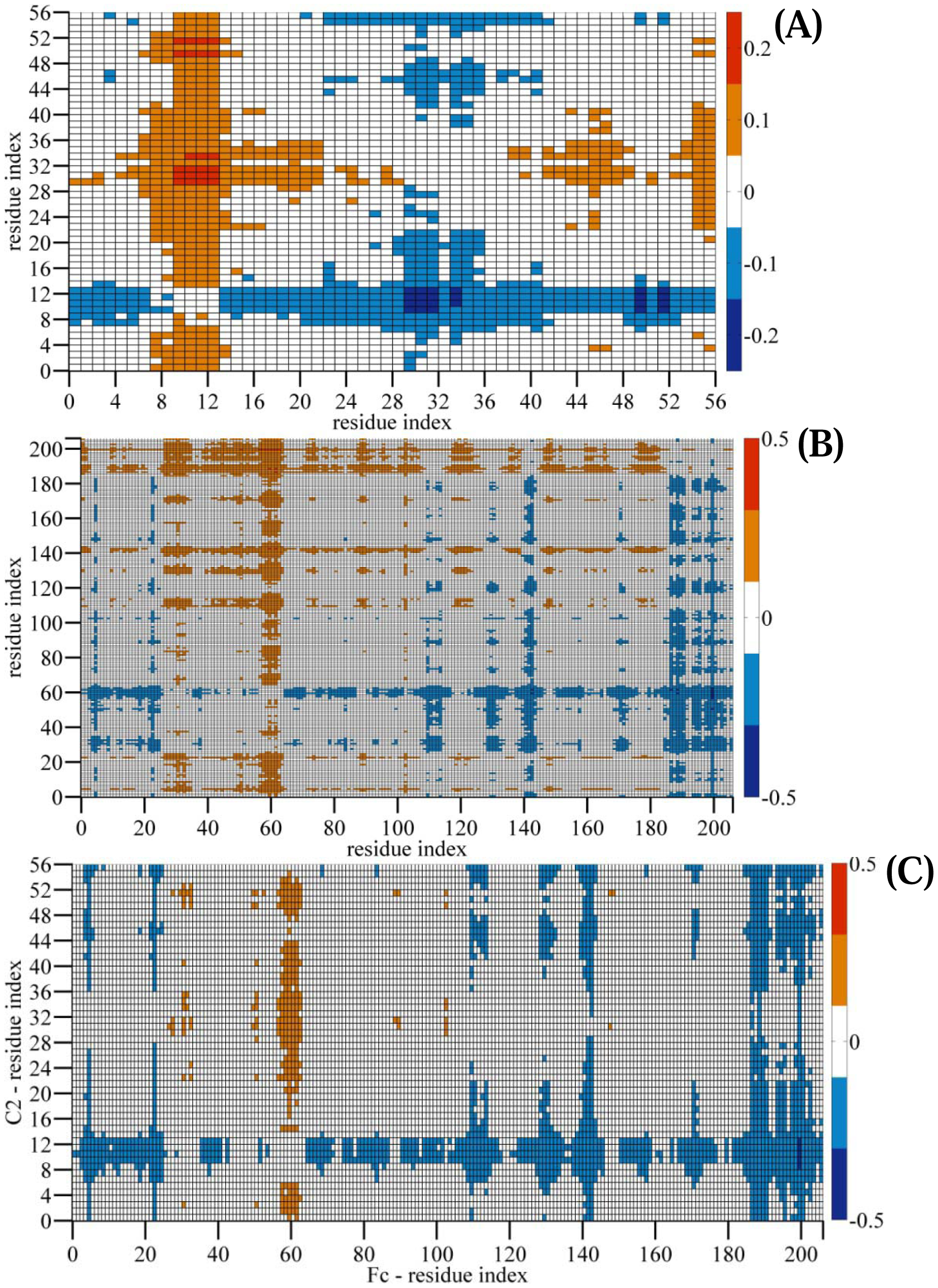
Directional Symbolic Transfer Entropy for the C2-Fc complex system: (A) Directional Symbolic Transfer Entropy of C2. (B) Directional Symbolic Transfer Entropy of Fc. (C) Directional Symbolic Transfer Entropy at the interface between C2 and Fc. Calculations are performed based on the average fluctuations of all backbone atoms for each residue. The trajectory lengths were 30 ns; 10000 snapshots were used to calculate the symbolic transfer entropy. Directional symbolic transfer entropy is presented as a color map using the scales shown next to the plots.

The scale of the values is shown in color bars plotted next to each graph. Our results identify the driving (source), characterized by a positive value of *D*_*i*→*j*_ and responding (sink) residues, characterized by a negative value of *D*_*i*→*j*_ That is, if *D*_*i*→*j*_ > 0, then residue *i* drives *j*, otherwise *j* drives *i*.

In Figure 15, we show the symbolic mutual information versus the time lag and false nearest neighbors count as a function of the embedded dimension *m*. It shows that mutual information drops exponentially with a time lag as expected. We determined the optimal time lag as the first minimum of the mutual information. Similarly, false nearest neighbors’ counts decrease with embedded dimension as shown in Figure 15B. We determined as the optimal value of embedded dimension the value for which the false nearest neighbors count becomes zero. From the inset graph in Figure 15B, the maximum value of *m* is around 6.

**Figure 15:**
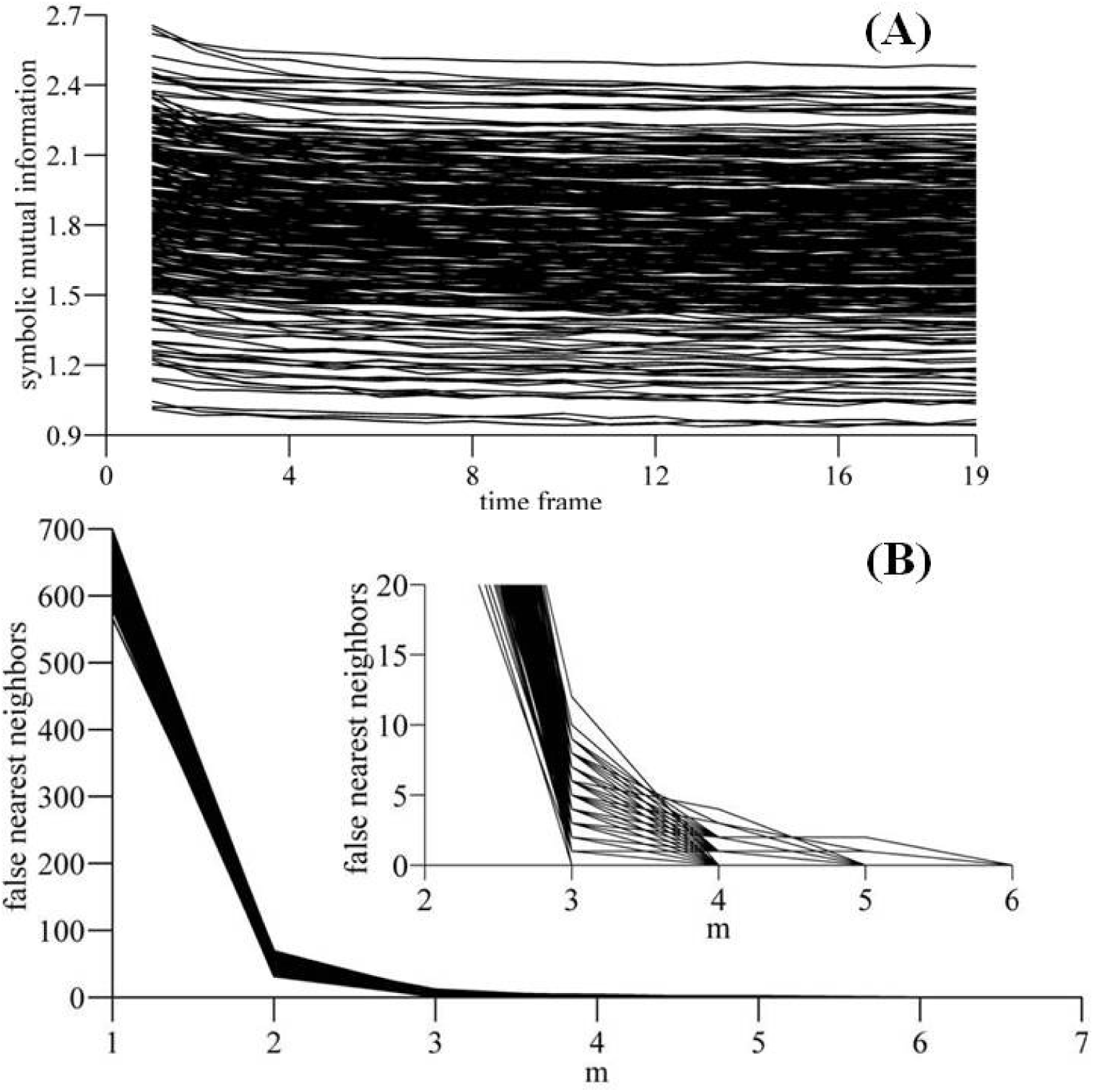
(A) Symbolic mutual information as a function of the time lag and (B) false nearest neighbors as a function of embedded dimension, *m*, for Protein-Protein complex system.

In Figure 16, we show the average values of the directional symbolic entropy for each residue for fragment C2 (Figure 16A) and Fc (Figure 16B). Besides, we show the distribution of the residues of the C2 fragment on the interface of the complex colored according to their average values of directional symbolic transfer entropy, namely in red a positive average value and in blue those with negative average values.

**Figure 16:**
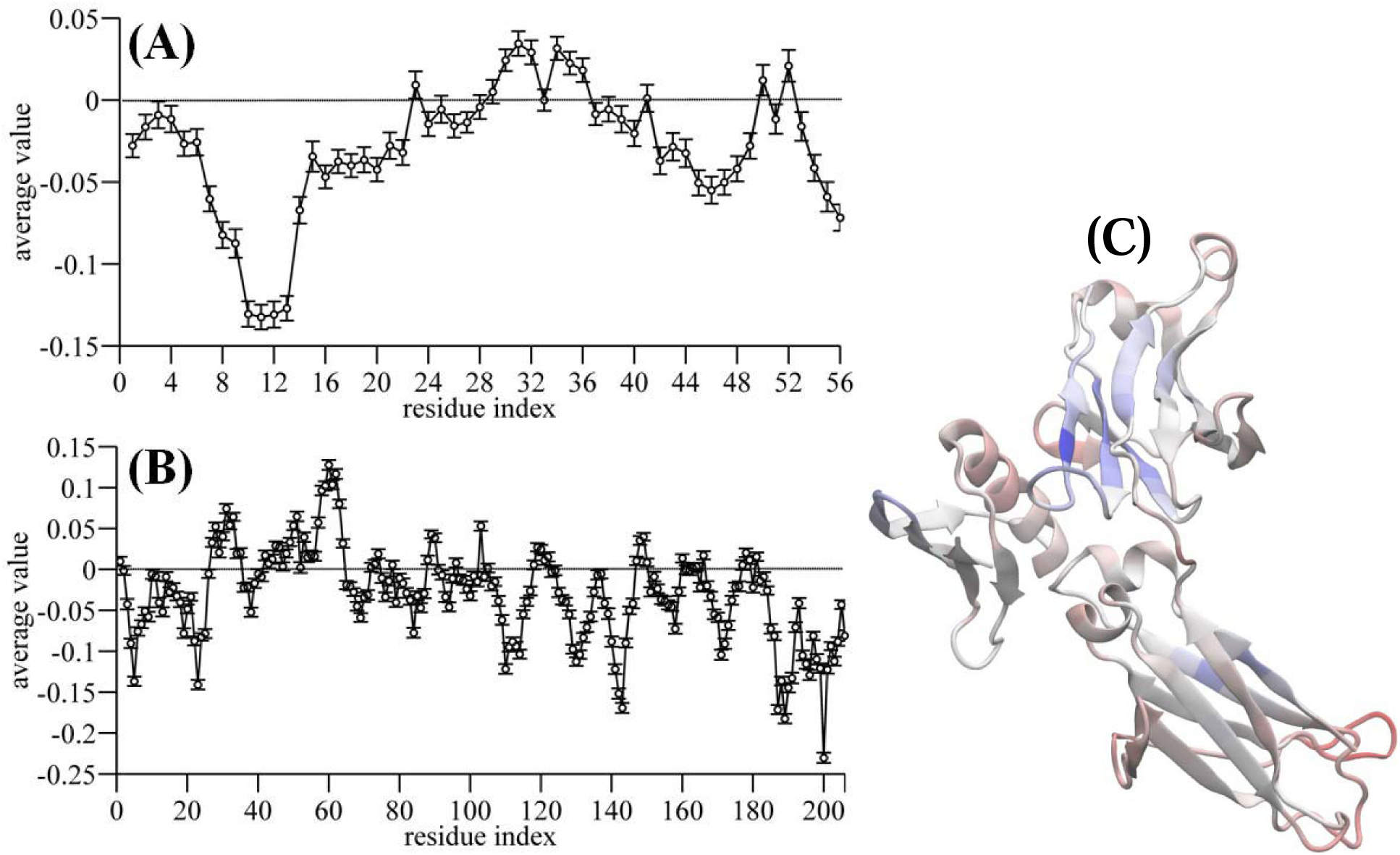
(A) The average directional symbolic entropy for each residue of the C2 fragment of protein G and (B) for Fc fragment of protein IgG. (C) A graphical representation of the amino acids colored according to their average values of the directional symbolic transfer entropy: in blue, negative values (*responding residues*) and in red, positive values (*driving residues*).

### Benchmark 3

Next benchmark test system is a complex biomolecular system, representing Protein – RNA interactions, as shown in Figure 17. The protein is composed of 88 amino acids, and RNA is composed of 6 bases.

**Figure 17:**
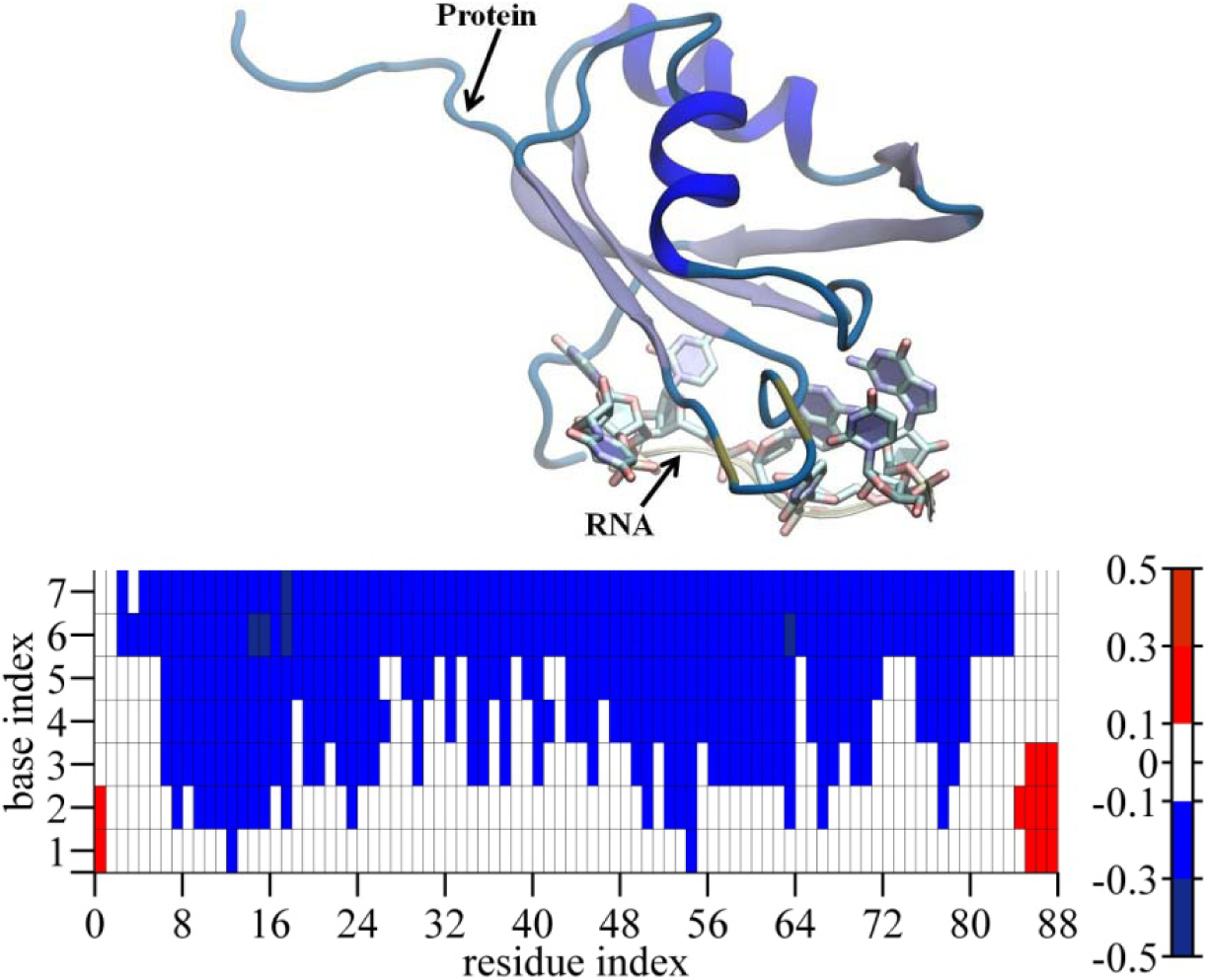
Complex biomolecular system representing a Protein-RNA complex system. Symbolic Mutual Information and Directional Transfer Entropy appear as color maps.

We calculated directional symbolic transfer entropy between protein and RNA, and we show it as a color map in Figure 17, along with a color bar indicating the scale of values. We can identify the residues driving the fluctuation motions and those responding to these fluctuations. In particular, we identify different clusters of residues acting as the source of the fluctuations: the first cluster includes the residues with the index from 1 to 2 and the second cluster includes the residues with the index from 7 to 12. There also exist three other small clusters driving the motions of the base 1 and 2, namely the cluster including the residues between 36 and 40, the cluster including the residues between 65 and 69 and this including the residues with an index between 75 and 87. In Figure 18, we show the averages of directional symbolic transfer entropy for each base and residue associated with all other residues or bases. Also, we present in blue the residues/bases acting as a sink and in red those acting as sources of the motion. It appears that they distribute along with the interface between protein and RNA.

**Figure 18:**
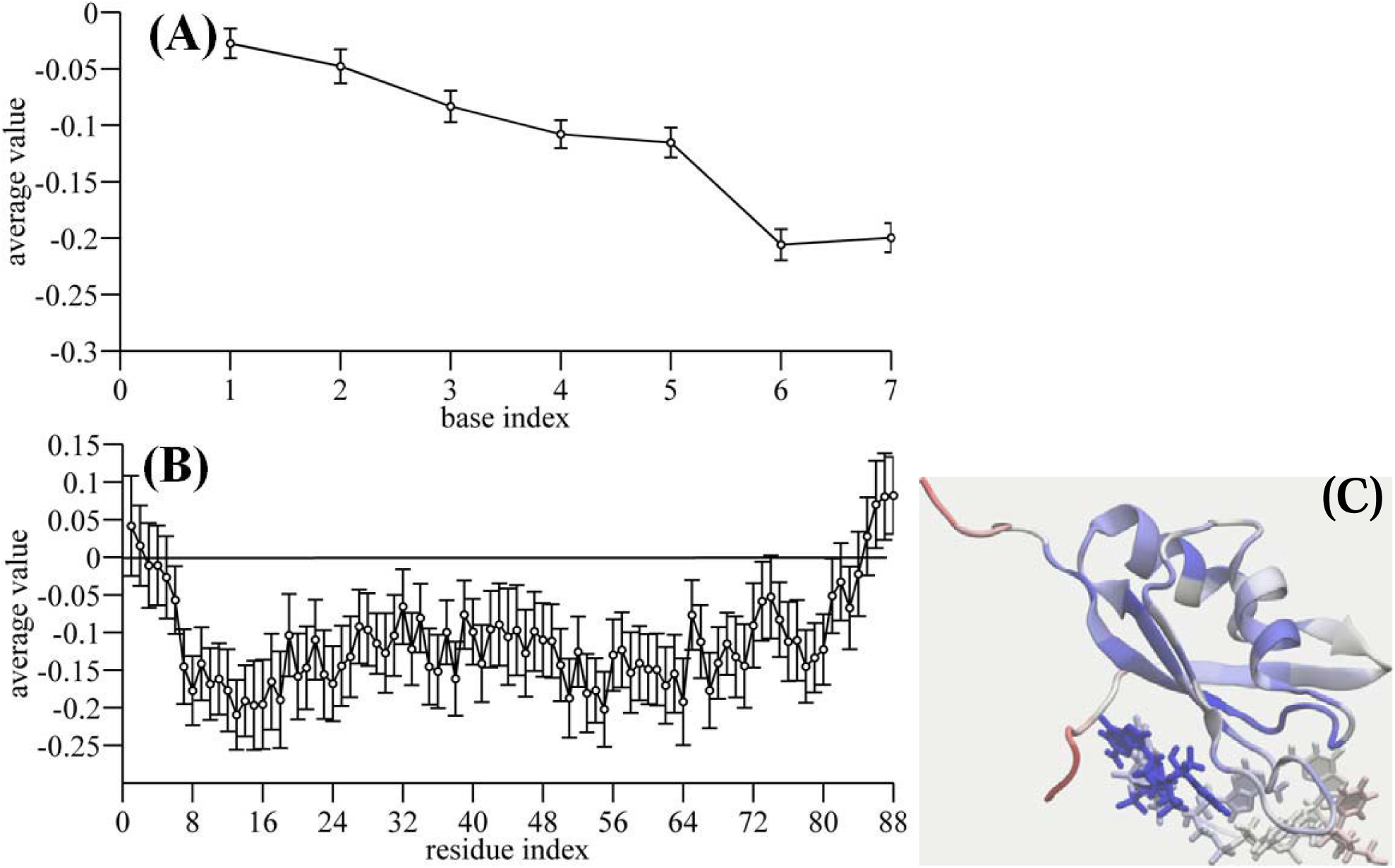
Complex biomolecular system representing a Protein-RNA complex system. Symbolic Mutual Information and Directional Transfer Entropy are shown as color maps.

Figure 19 shows the symbolic mutual information, calculated according to Eq. (26), as a function of the time lag and false nearest neighbors as a function of the embedded dimension. The results are used to calculate both time lag *τ* and embedded dimension *m.* Clearly, our data show that symbolic mutual information decreases exponentially with a time lag, and here we considered as the optimal time lag the value at with mutual information reaches the first minimum. Furthermore, the false nearest neighbors values show a fast decrease with *m* to zero, which is reached for *m* between 4 and 6, as indicated from the inset graph in Figure 19B.

**Figure 19:**
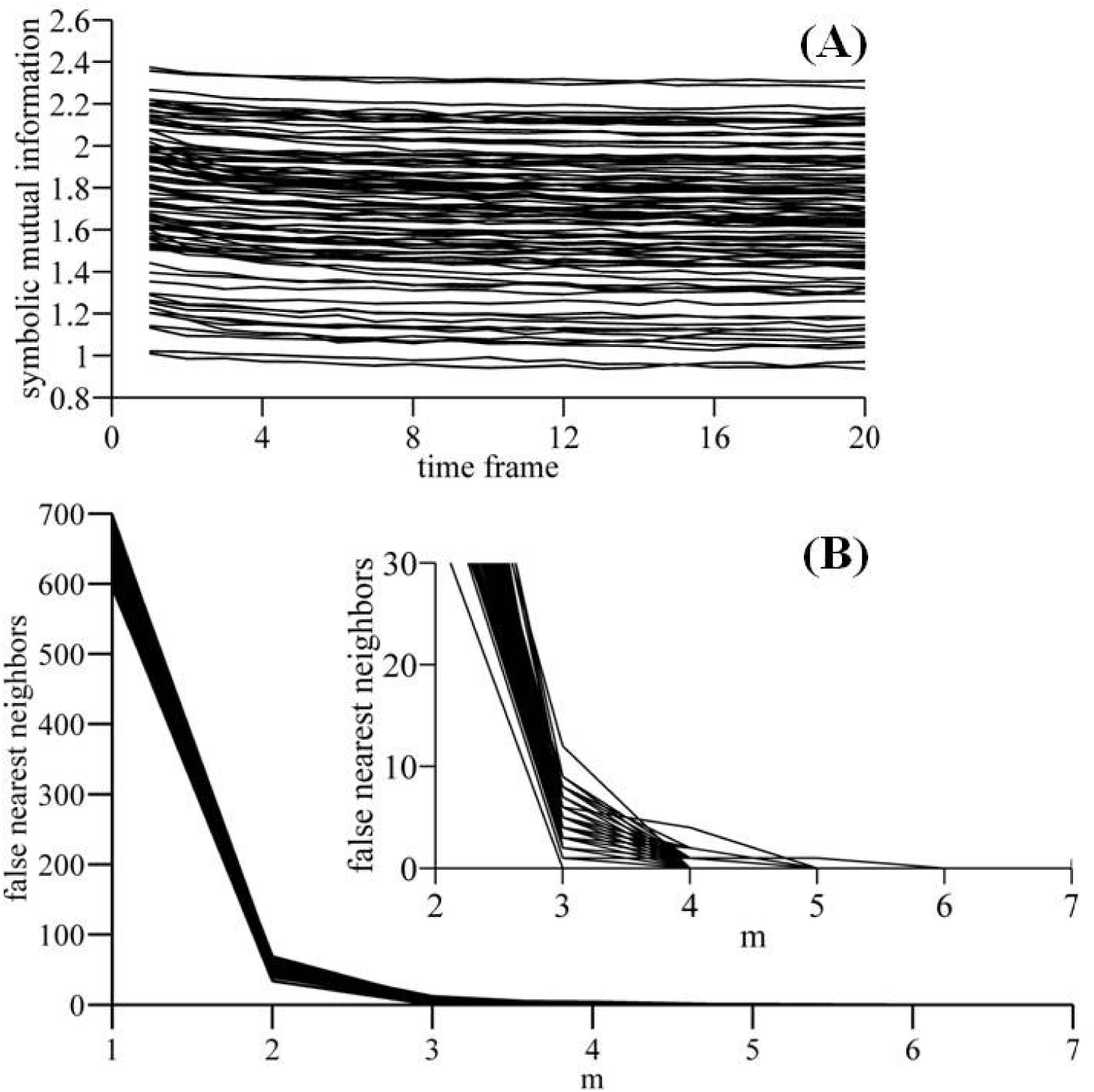
(A) Symbolic mutual information as a function of the time lag and (B) false nearest neighbors as a function of embedded dimension, *m*, for Protein-RNA complex system.

## 6 Conclusions

In this study, we present new software for computation of symbolic analysis of time series representing random processes of dynamical systems. Besides, the software can compute symbolic transfer entropy, symbolic mutual information, and symbolic local transfer entropy, where definitions of the symbolic mutual information and symbolic transfer entropy to the best of our knowledge are given for the first time in this study.

Also, we present a new method using a machine learning approach on how to reduce the dimensionality of the dynamical variables characterizing the different systems or different components of a dynamical system. We envision that the algorithm presented in this study can be used to measure the information flow between components of a dynamical system and to identify the direction of the information flow between different dynamical systems, such as in the signal pathways problems.

Some of the functionalities of this software added during the implementation include portability, adjustable computational precision, efficient memory management, efficient and practical data management system, parallelization of computations using MPI protocols, and graphical user interface.

## * Acknowledgements

Authors like to thank the International Balkan University for support.

## B Required Metadata

### B1 Current executable software version

**Table 1.**
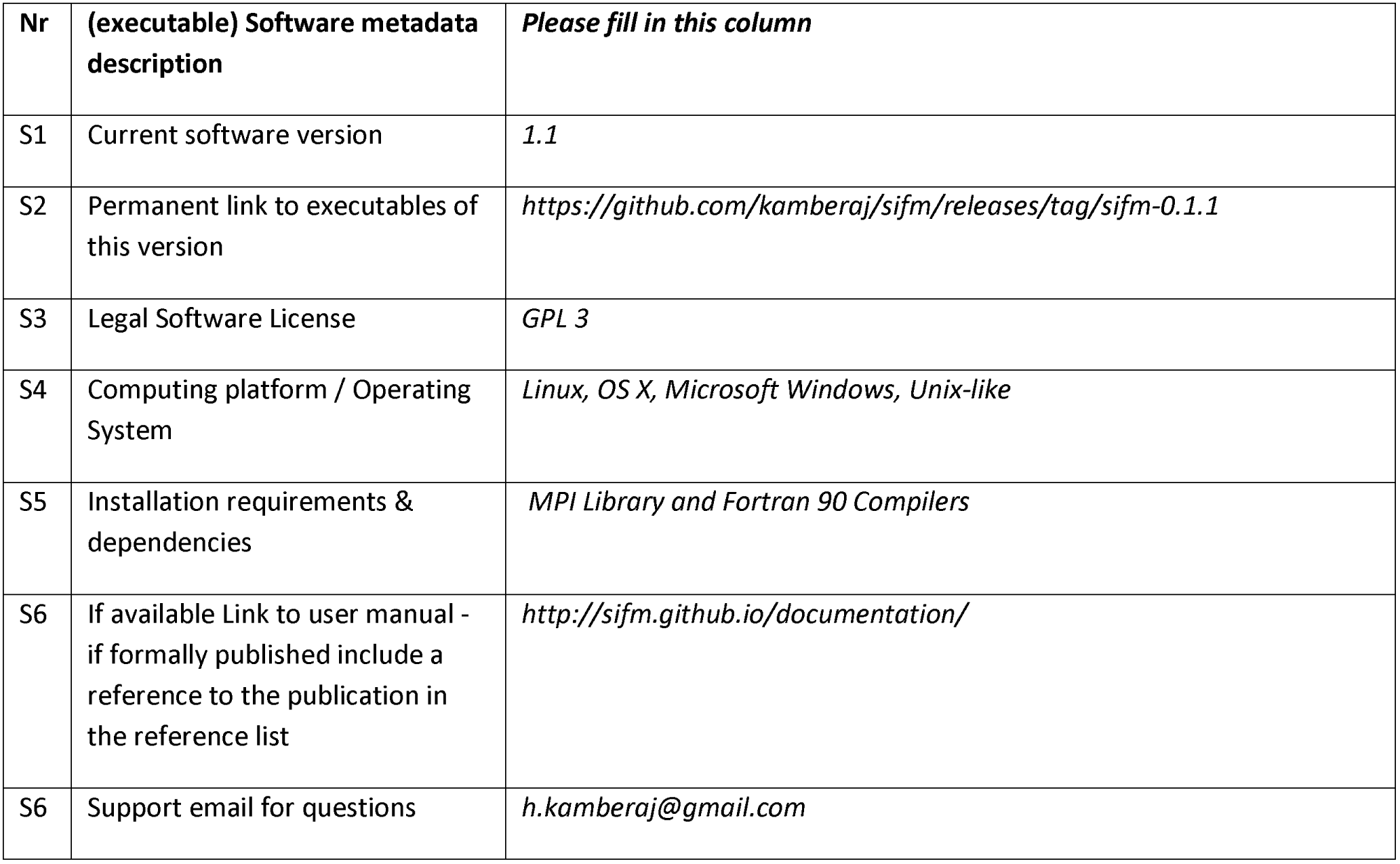
Software metadata.

### B2 Current code version

**Table 2.**
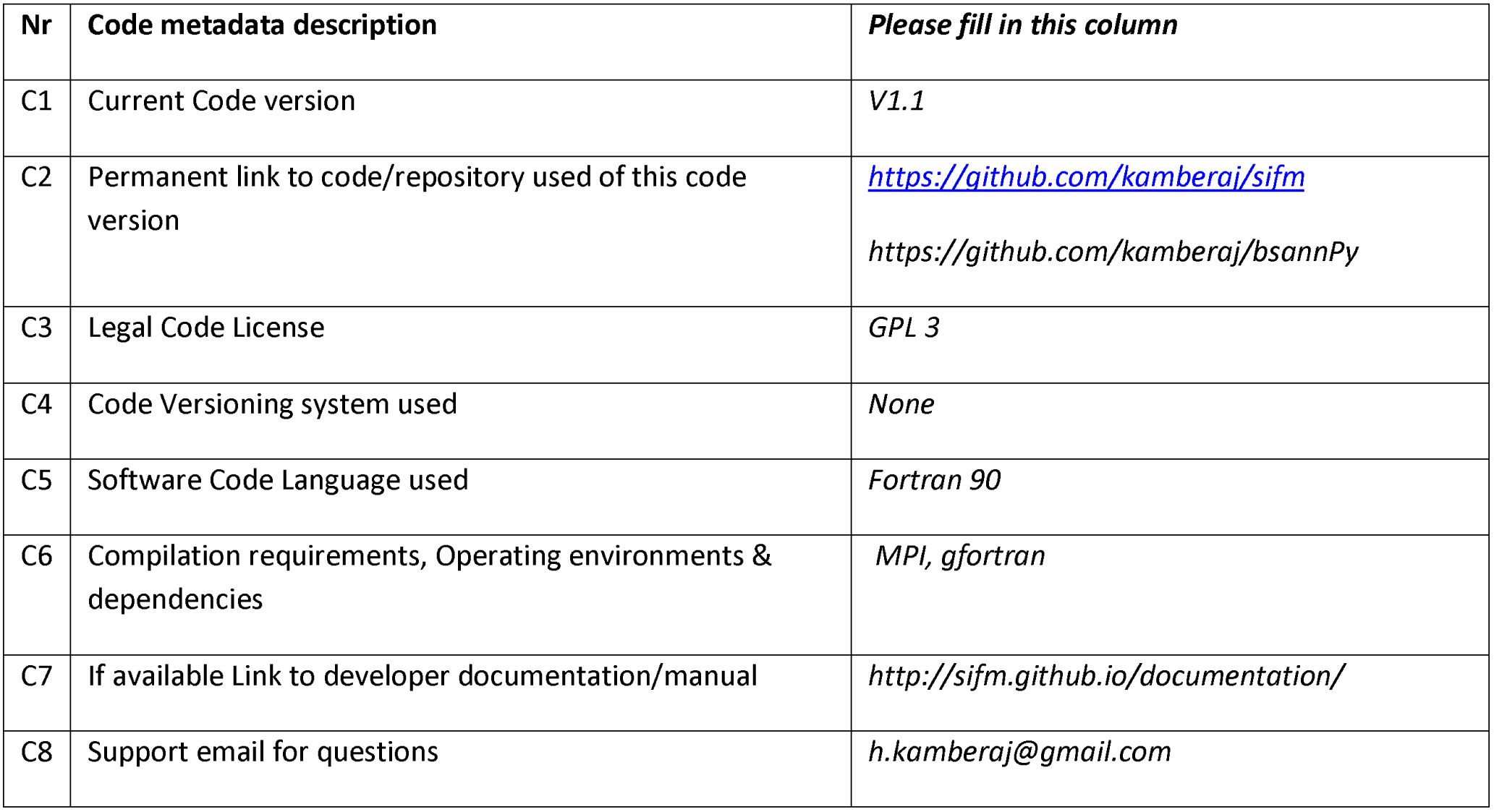
Code metadata.

